# Targeting PD-1^+^ T-cells with Chimeric Antigen Receptors to reduce the HIV Reservoir

**DOI:** 10.1101/2025.09.10.675382

**Authors:** Laura Ermellino, Riddhima Banga, Spiros Georgakis, Nicole P. Kadzioch, Francesco Procopio, Ana Alcaraz-Serna, Oscar Alfageme-Abello, Raphaël Porret, Rebecca Cecchin, Michail Orfanakis, Rachel Schelling, Cloé Brenna, Duy-Cat Can, Mathilde Foglierini, Oliver Y. Chén, Laurent Perez, Craig Fenwick, Matthieu Perreau, Constantinos Petrovas, Roberto F. Speck, Giuseppe Pantaleo, Yannick D. Muller

**Author notes:** Contributed equally, authors shown in alphabetic order (Riddhima Banga, Spiros Georgakis, Nicole P. Kadzioch, Francesco Procopio). <u>Corresponding author:</u> Yannick Daniel Muller – MD-PhD – Assistant Professor, Service d’immunologie et d’allergie, Département de médecine, BH010-511, Rue du Bugnon 46, CH-1011 Lausanne.

## Abstract

The unique ability of chimeric antigen receptor (CAR) T-cells to infiltrate tissues is revolutionizing our perspectives for tackling severe-refractory and otherwise untreatable diseases. In HIV, CAR-T-cells have been designed to target viral biomarkers, with limited success so far. Here, we investigated the possibility of redirecting CAR-T-cells against a cellular biomarker of the HIV reservoir, PD-1. We designed two second-generation 4-1BB-CARs using the scFv of either a blocking (bPD1-CAR) or a nonblocking (nbPD1-CAR) anti-PD-1 monoclonal antibody. The CAR avidity modulated T-cell sensitivity, trogocytosis, and effector functions, independently of the PD-1 signalling domain. Both anti-PD-1 CAR T-cells could persist for 70 days in HIV-infected humanized mice, correlating with viral protection and a disruption of the lymphoid architecture in the white pulp of the spleen. Altogether, our results open new strategic avenues for reducing the HIV reservoir as we demonstrate the feasibility of depleting specific T-cell subpopulations.

**Summary:** T cells can be redirected against cellular rather than viral-specific biomarkers to reduce the HIV reservoir.

## INTRODUCTION

Chimeric Antigen Receptor (CAR) T cells are revolutionizing strategic therapeutic approaches for tackling severe, refractory, and otherwise untreatable diseases. Beyond the success of CAR T cells in oncological conditions (*1*), the past years have seen the emergence of cellular approaches to treat systemic lupus erythematosus, idiopathic inflammatory myositis, systemic sclerosis, but also neurological and other autoimmune conditions (*2*–*4*). The unexpected results of long-term remission in patients who otherwise require chronic immunosuppression have led to the concept of immune resetting (*5*). Thus, CAR-T cells possess unique abilities to clear the autoreactive B cell repertoire within tissues and lymphoid organs, where monoclonal antibody-based therapies (mAb) have failed (*6*).

The first clinical trials using engineered T cells were done in the field of HIV in the early nineties. Conceptually, T cells were redirected against the HIV envelope, taking advantage that HIV-infected cells can be detected because they express viral-specific biomarkers at their cell surface (*7*). Thus, T cells have been first engineered with a CD4 extracellular domain fused to a CD3ζ signaling domain to kill HIV-infected cells (*8*, *9*). Since the emergence of broadly neutralizing antibodies (*10*, *11*), T cells have been armed with single chain variable fragments (scFv) targeting several regions of the HIV envelope in addition to the CD4 binding site such as the V1V2 apex or the membrane-proximal external region (MPER) of gp41 glycan site. Although no benefit has been reported in CAR-T treated HIV-infected patients to date, several clinical trials are ongoing evaluating dual/bi-specific anti-HIV CARs (*12*).

Over the last two decades, much effort has been employed to understand the complexity of the HIV reservoir in order to foster the development of therapeutic interventions that could lead to a functional cure of HIV. Several subsets of CD4^+^ T cells harbor transcriptionally silent proviruses at different frequencies and can escape immune surveillance (*13*). In particular, there is a significant viral enrichment within the lymph nodes of aviremic individuals treated with antiretroviral therapies (ART) in CD4^+^ T cells expressing PD-1, namely follicular helper cells (Tfh) (*14*, *15*). Others found elevated levels of integrated HIV DNA in circulating memory PD-1^+^ CD4^+^ T cells (*15*, *16*). Finally, PD-1 has also been described as a marker for HIV persistence in other anatomical sites such as rectal tissues (*17*).

Considering the unique abilities of T cells to migrate deep into the tissues, we hypothesize that T cells can be redirected against the cellular marker PD-1, which could contribute in reducing the HIV reservoir. We developed second-generation 4-1BB CARs using scFv derived from two anti-PD-1 antibodies, one competing with PDL-1 (blocking (b)PD1-CAR) and the second targeting a non-competitive allosteric epitope of PD-1 (non-blocking (nb)PD1-CAR) (*18*, *19*). We showed that the functional activity of engineered T cells is modulated by the CAR avidity. Both CAR-T cells were detected following adoptive cell transfer (ACT) correlating with reduced number of PD-1^+^ cells, delayed HIV rebound and loss of the lymphoid architecture in the white pulp of the spleen of humanized mice. These findings underscore the potential of anti-PD-1 CAR-T cells in resetting the lymphoid architecture and thereby reducing the size of the HIV reservoir.

## RESULTS

### Blocking and non-blocking anti-PD-1 antibodies characterization and CAR design

We selected the scFv of two anti-PD-1 monoclonal antibodies previously discovered by our center, interacting with two different regions of the PD-1 receptor: clone A35795 (A35) blocking the interaction with PD-L1 and clone 135c139d6 (135C) binding to a different epitope independently of the PD-1/PD-L1 engagement site (Fig.1A) (*18*, *19*). We produced IgG1 antibodies and confirmed their binding to Jurkat cells overexpressing PD-1 (Fig. 1B). A competitive binding assay with increasing concentration of anti-PD-1 monoclonal antibodies Pembrolizumab or Nivolumab demonstrated the blocking and non-blocking nature of clones A35 and 135C respectively (Fig. 1B and Fig. S1A). To further evaluate the affinity of both clones, we produced them as fragment antigen-binding domains (Fab) and single chain variable fragments (scFv). Both Fabs had a similar dissociation constant (KD) in the range of 10-16 nM, while clone 135C, when expressed as scFv, exhibited a four times lower KD than clone A35 (Fig. 1C and Fig. 1D). We next engineered human T cells expressing both scFvs as a 4-1BB second-generation CAR flanked to a ribosomal skipping motif and mCherry reporter. In contrast to the results obtained with purified scFv, the binding of biotinylated PD-1 was lower for the 135C CAR suggesting a lower avidity (Fig. 1E and Fig. S1B).

**Fig. 1.**
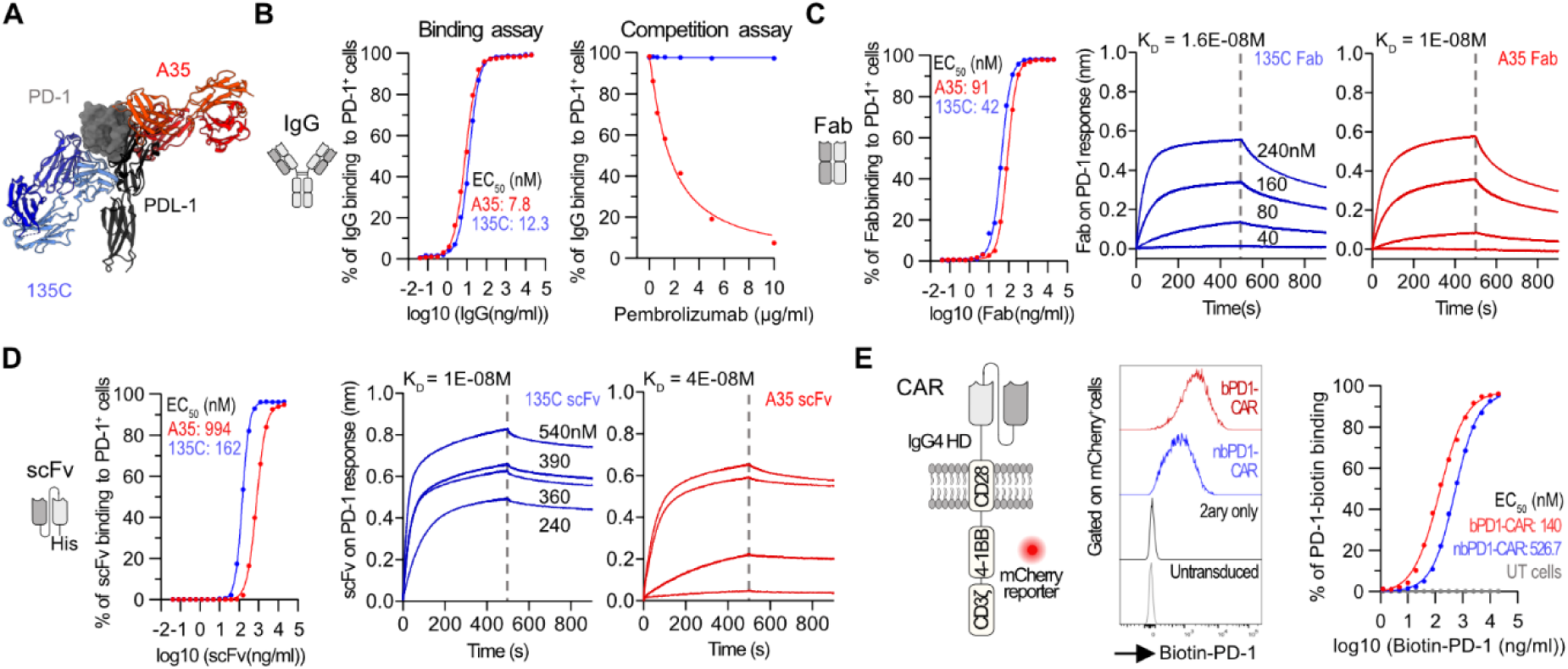
PD-1 binding to CAR-T cells is epitope dependent and not necessarily predicted by the antibody affinity. (A) In silico model representing the binding of Fab fragment from clones A35795 (A35) and 135c139d6 (135c) compared to the hPD-1/hPD-L1 complex (PDB ID: 4ZQK). (B) Binding of A35 and 135c IgG to human PD-1 (left). Competitive binding assay with Pembrolizumab and biotin-labelled A35 IgG or 135c IgG (right). Symbols are means of 3 (left) or 2 (right) independent experiments. (C) Binding of A35 and 135c Fabs to human PD-1 (left). Biolayer interferometry of Fabs–PD-1 interactions (right). Symbols are means of 3 independent experiments (left). (D) Binding of A35 and 135c scFvs-His to human PD-1 (left). Biolayer interferometry of scFvs–PD-1 interactions (right). Symbols are the mean of 3 independent experiments (left). (E) Second-generation CAR design (left). Biotinylated PD-1 binding to anti-PD-1 CARs. Representative flow cytometry and mean of 4 independent experiments. <u>Abbreviations</u>. PD-1, Programmed Cell Death Protein 1; PD-L1, Programmed Cell Death Ligand 1; IgG, Immunoglobulin G; Fab, Fragment Antigen-binding region; K_D_, equilibrium dissociation constant; scFv, single chain variable fragment; His, Histidine; CAR, Chimeric Antigen Receptor; EGFRt, truncated Epidermal Growth Factor Receptor; HD, hinge domain; bPD1-CAR, blocking anti-PD-1 CAR; nbPD1-CAR, non-blocking anti-PD-1 CAR; UT, untransduced cells.

### Anti-PD-1 CARs cytotoxic activity is single-chain variable fragment dependent

To further evaluate the functional activity of both CARs in primary T cells, we included an anti-CD19 CAR control in addition to the non-blocking (nbPD1, 135C) and blocking (bPD1, A35) CARs. All CARs had a Myc tag at the N-terminus and expressed mCherry as a reporter (Fig. 2A). We obtained comparable surface expression, transduction efficiencies and expansion rates of all three CARs (Fig. 2B and Fig. S2A, B). Notably, we found a spontaneous and significant loss of PD-1^+^ cells, mainly in the Cherry^-^ population of the bPD1-CAR condition (Fig. 2D and Fig. S3A). Such depletion was also present in the nbPD1-CAR condition but only at a high transduction level (Fig. 2D and Fig. S3A). Interestingly, the PD-1 loss found with the bPD1-CAR correlated with a significant reduction in the number of CD4^+^ T cells, likely because CD8^+^ T cells expressed lower levels of PD-1 during in vitro expansion (Fig. 2E, Fig. S3B). Importantly, when performing a knockout of PD-1 (day 0) prior to CAR editing (day 1), we could prevent the loss of CD4^+^ T cells (Fig. 2E and Fig. S3C), confirming the hypothesis that CD4^+^ T cells loss was related to their PD-1 surface expression.

**Fig. 2.**
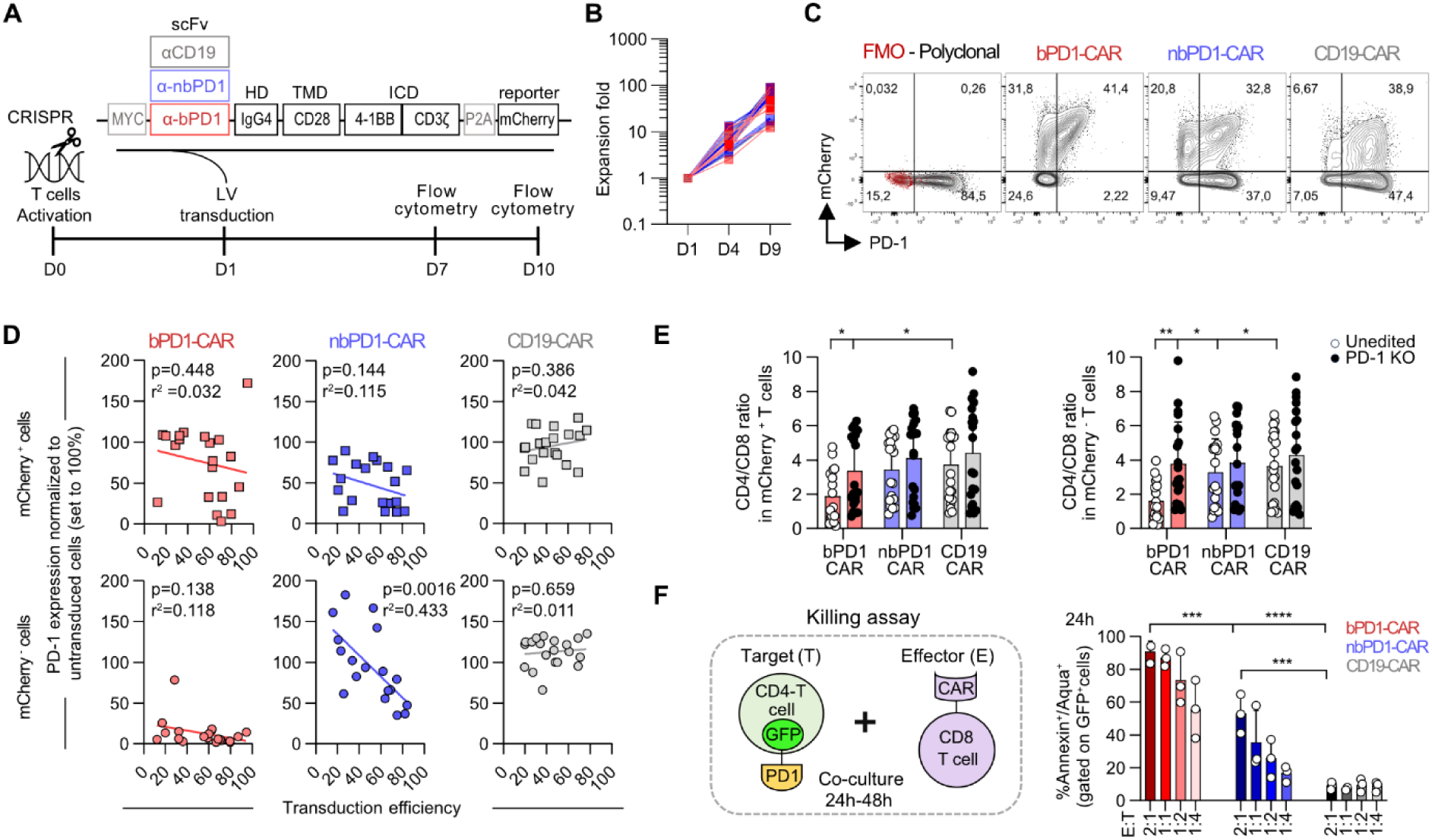
The non-blocking anti-PD-1 CAR showed intermediate functional activities in vitro. (A) Experimental design for engineering human anti-PD-1 CAR-T cells. (B) Expansion of primary anti-PD-1 CAR-T cells compared to the anti-CD19 CAR control (n=7 donor, 7 independent experiments). (C) Representative flow cytometry plots showing mCherry and PD-1 expression 6 days after transduction (gated on CD3^+^ living cells). (D) Correlation between PD-1 expression and the percentage of transduction efficiency. Cherry positive and negative population are shown. For each experiment the percentage of PD-1+ cells were assessed in Cherry negative and positive cells and normalized to PD-1 expression in untransduced T cells (set as 100%). 8 donors, 8 independent experiments each with 2-3 internal replicates (with different MOI transduction:1.4 ± 0.6). Simple linear regression tests. (E) CD8/CD4 ratio in mCherry +/- T cells comparing anti-PD-1 CAR and CD19 CAR-T cells +/- PD-1 knockout. Data are presented as Mean ± SD values of 8 donors, 8 independent experiments, 2-3 internal replicates). 2way ANOVA, Tukey’s multiple comparisons test. (F) Schematic representing the killing assay (left). Cumulative percentage of Annexin/Aqua^+^ CD4^+^PD-1^+^GFP^+^ target cells after 24h of co-culture (right). Mean ± SD of 3 donors and 3 independent experiments is shown. Two-way ANOVA, Tukey’s multiple comparison test. <u>Abbreviations</u>. CRISPR, Clustered Regularly Interspaced Short Palindromic Repeats. HD, Hinge Domain. TMD, Trans-Membrane Domain. ICD, Intracellular Domain. LV, lentivirus. scFv, single chain variable fragment. CAR, Chimeric Antigen Receptor. bPD1-CAR, blocking anti-PD-1 CAR. nbPD1-CAR, non-blocking anti-PD-1 CAR. FMO, Fluorescence minus one. UT, untransduced. KO, knock out.

To evaluate the cytotoxicity of both CARs, we next expanded separately human CD8^+^ CAR-T cells from CD4^+^ T cells which were transduced with a PD-1-GFP fusion protein to ensure stable PD-1 expression (Fig. S4A, B). The CD4^+^ T cells were killed by both CARs, the bPD1-CAR being more efficient (Fig. S4C, D and Fig. 2F).

### bPD1-CAR-T cells show enhanced sensitivity and cytotoxicity independently of the PD-1 signaling domain

To better characterize the functional activity of anti-PD-1 CARs, we next assessed their sensitivity and PD-1 trogocytosis capacities. Starting from a luciferase positive PD-1 KO Jurkat cell line, we generated seven clones with increasing levels of PD-1 ranging from 424 to 6047 PD-1 molecules and used them as targets in a bioluminescent killing assay. To further reduce the potential off-target activities we co-edited TRBC (T cell receptor beta chain) in addition to the endogenous PD-1 receptor of primary T cells (Fig. S5A-C). Consistently with the in vitro killing results in primary cells, bPD1-CAR T cells exhibited higher sensitivity with low expressing cell lines as well (Fig. 3B). For the trogocytosis assay, we transduced K-562 PD-1 KO cells with a lentiviral vector coding for GFP-only or a PD-1-GFP fusion protein (Fig. 3C). In line with the sensitivity results, the bPD1-CAR acquired GFP-PD-1 more efficiently than the nbPD1-CAR (Fig. 3D, E).

**Fig. 3.**
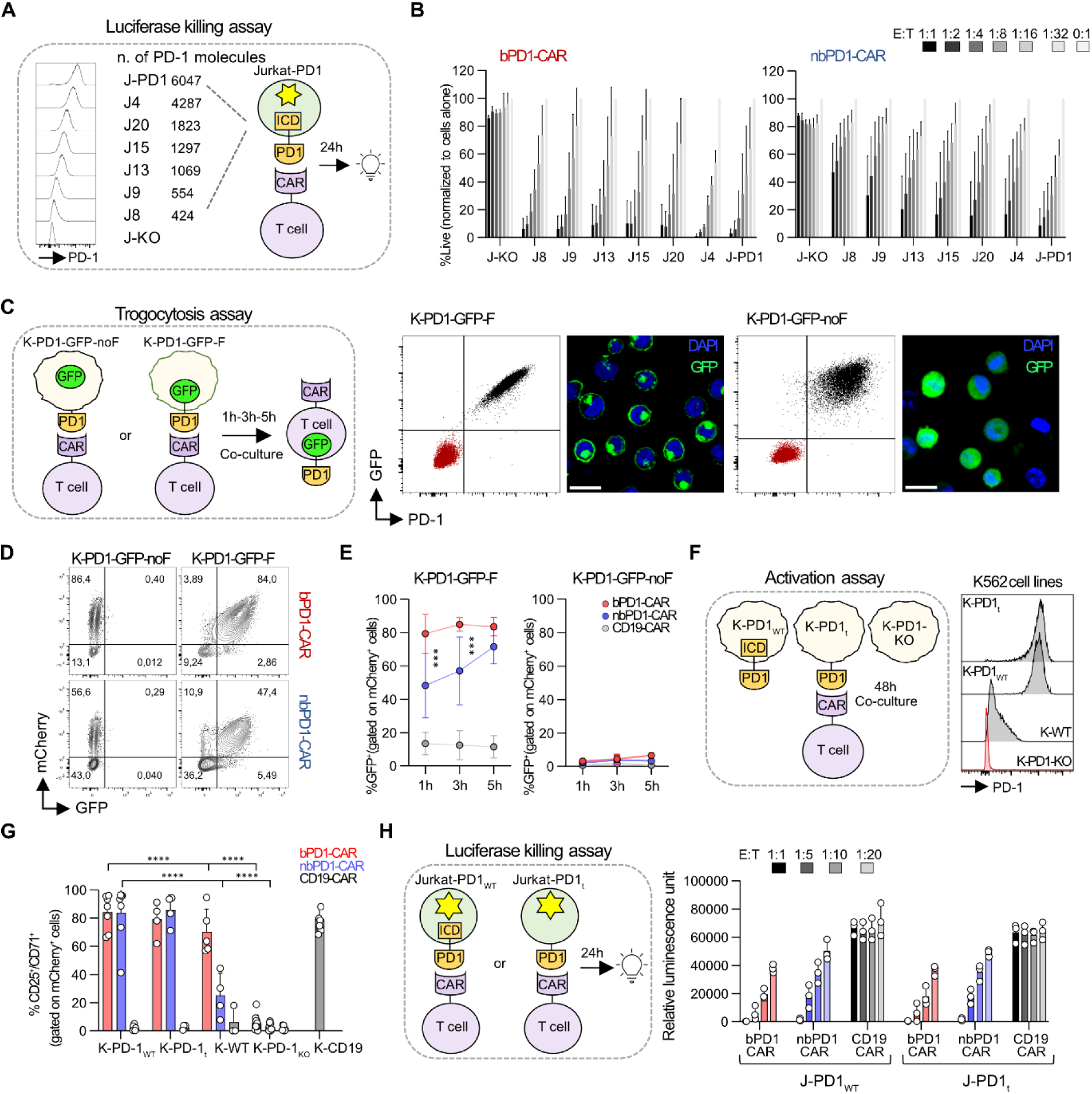
PD-1 trogocytosis is CAR dependent and does not inhibit T cells activation. (A) Generation of PD-1-transgenic luciferase^+^ Jurkat cells with increasing PD-1 molecules number. (B) Luciferase-based killing assay of blocking and non-blocking CAR-T cells. Mean ± SD of 3 donors and 3 independent experiments is shown. (C) Phenotype of the K562 cell lines expressing either a PD-1-GFP-F fusion protein (K-PD-1-GFP-F) or PD-1 and GFP proteins with a ribosomal skipping motif in-between (K-PD-1-GFP-noF) used for the trogocytosis assay. Scale bar is 25 μm. (D) Representative flow cytometry 1h after co-culture of K-PD-1-GFP-F or -PD-1-GFP-noF with anti-PD-1 CAR-T cells. (E) Cumulative data showing GFP expression in CAR-T cells over time. Mean ± SD values of 6 donors and 4 independent experiments is shown. Two-way ANOVA, Tukey’s multiple comparison test. (F) Phenotype of the K562 cell lines expressing either the wild type PD-1 (K-PD-1_WT_) or a truncated PD-1 lacking (K-PD-1t) the intracellular domain used for the activation assay. (G) Cumulative percentage of CD25^+^/CD71^+^ CAR-T cells after a 48h co-culture with K-PD-1_WT_ or K-PD-1t. Mean ± SD values of 4-8 donors and 7 independent experiments is shown. Two-way ANOVA, Tukey’s multiple comparison test. (H) Luciferase killing assay using PD-1_WT_ or PD-1t luciferase^+^ Jurkat cells as targets. Mean ± SD values of 3 donors, 3 independent experiments is shown. Two-way ANOVA, Tukey’s multiple comparison test was used to compare the same condition in JPD-1 vs JPD-1_t_. <u>Abbreviations</u>. JPD-1, Jurkat-PD-1. ICD, Intracellular Domain. CAR, Chimeric Antigen Receptor. bPD1-CAR, blocking anti-PD-1 CAR. nbPD1-CAR, non-blocking anti-PD-1 CAR. E:T, Effector: target. K-PD-1-GFP-F, K562 PD-1-GFP-F fusion protein. K-PD-1-GFP-noF, K562 PD-1+ GFP+. K-PD-1, K562 PD-1+. K-PD-1-t, K562 PD-1 truncated. K-PD-1-KO, K562 PD-1 KO. JPD-1-t, Jurkat PD-1 truncated.

Given that the intracellular domain (ICD) of PD-1 mediates inhibitory signaling and the CARs ability to uptake PD-1, we next evaluated the contribution of the PD-1-ICD to CAR-T cell functional activity. We engineered three K562 cell lines expressing PD-1 either with a truncated ICD (PD-1t) or the wild type molecule (PD-1_WT_) and compared them to a WT K562 cell line spontaneously expressing low levels of PD-1 (Fig. 3F). Importantly, we did not observe any differences in the level of activation or in the cytotoxicity between the truncated and WT PD-1 for both PD-1^high^ transgenic cell lines, suggesting that the uptake of PD-1 has no direct impact on CAR-T cell function (Fig. 3G, H, Fig. S5D).

### T cells exhaustion and terminal differentiation are anti-PD-1 CAR dependent

Since anti-PD-1 CAR-T cells naturally upregulate PD-1 upon activation and can uptake external PD-1 by trogocytosis, we next evaluated whether this could contribute to repetitive stimulation ultimately accelerating T cell differentiation and exhaustion. As a positive control, we engineered HLA-A2^+^ T cells with an anti-HLA-A2 (A2) CAR using a previously reported scFv (*20*), and as negative controls, we deleted either HLA-A2 or PD-1 from the A2 or PD1-CAR-T cells respectively (Fig. 4A, Fig. S6A, B). While for the bPD1 CAR condition, PD-1 expression was strongly reduced, for the HLA-A2 CAR condition, HLA-A2 remained highly expressed in both cherry positive and negative cells (Fig. 4B, C). Consequently, HLA-A2 CAR-T showed significantly reduced expansion and a terminally differentiated profile, as indicated by the expression of exhaustion makers (Lag3/Tim3). This phenotype was restored by editing HLA-A2 (Fig. 4D-F, Fig. S6C-F). To a lesser extent, the bPD1-CAR-T cells also showed higher levels of exhaustion and terminal differentiation, which interestingly was not the case for the nbPD1-CAR condition. Altogether, these results suggest that the inducible nature of PD-1 can preserve anti-PD-1 CAR-T cells from early exhaustion in a CAR-avidity dependent manner.

**Fig. 4.**
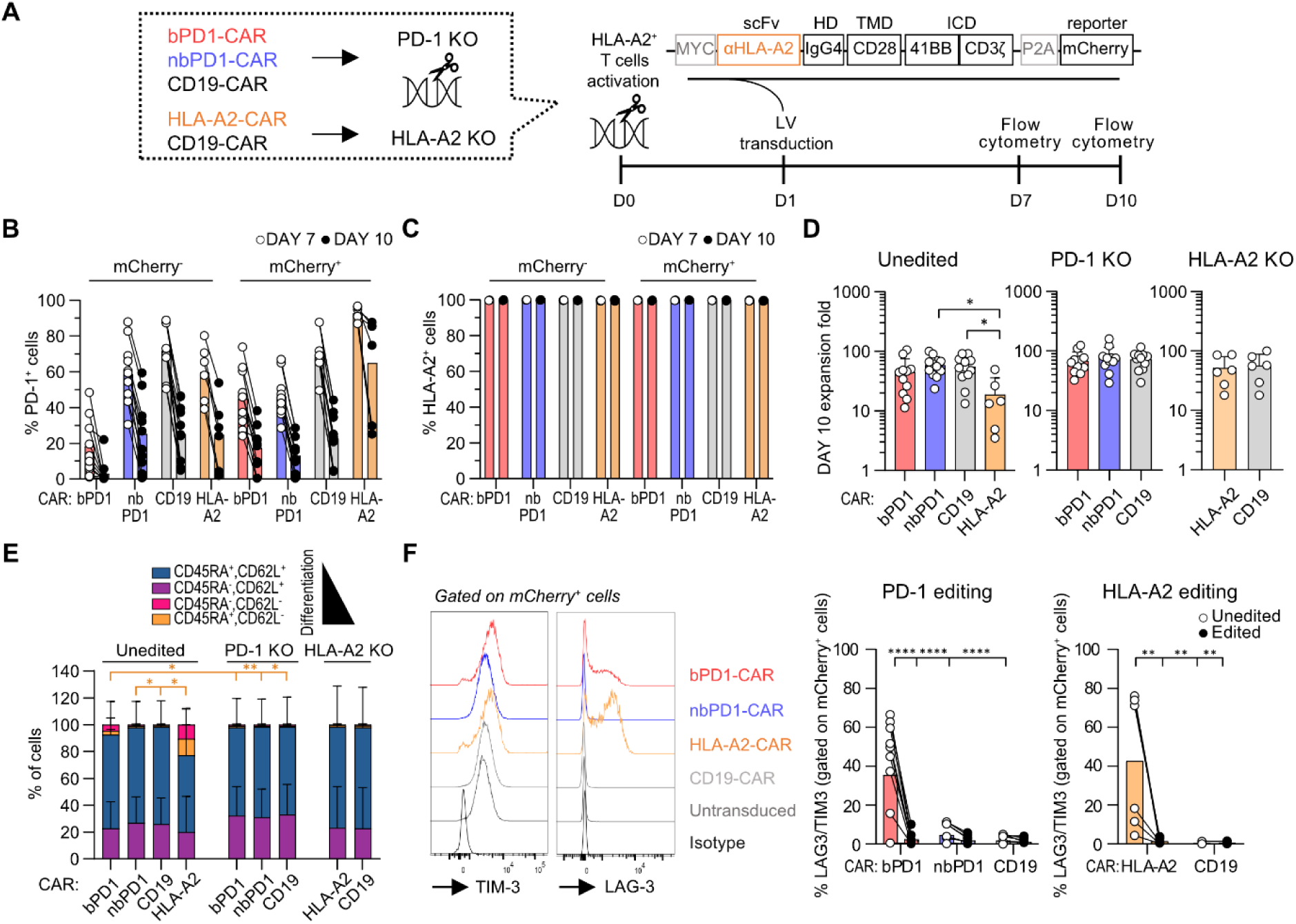
Constitutive expression of endogenous receptors and CAR-T cells sensitivity modulate terminal differentiation, exhaustion and cell death. (A) Experimental design. HLA-A2+ human T cells were isolated and activated +/- prior editing of PD-1 and/or HLA-A2. (B) PD-1 expression in mCherry^+^ and mCherry^-^ cells on day 7 and 10. Mean of 3-5 donors in 3-5 independent experiments with 1-2 replicates transduced at different MOI (1.4 ± 0.6) for each independent experiments. (C) HLA-A2 expression in mCherry^+^ and mCherry^-^ cells on day 7 and 10. Mean of 2 donors in 2 independent experiments with 1-2 internal replicates (different MOI transduction:1.4 ± 0.6). (D) Expansion folds of unedited versus edited CAR-T cells on day 10. Mean ± SD values of 3-5 donors in 3-5 independent experiments with 1-2 internal replicates (different MOI transduction: 1.4 ± 0.6). Statistics: Ordinary one-way ANOVA, Tukey’s multiple comparison test, for Unedited and PD-1 KO condition. Two-tailed paired T test for HLA-A2 KO condition. (E) Cumulative data showing CD62L/CD45RA expression of mCherry^+^ cells in edited versus unedited condition for PD-1 or HLA-A2. The mean ± SD values of 3-5 donors in 3-5 independent experiments with 1-2 internal replicates (different MOI transduction 1.4 ± 0.6) are shown. Kruskal-Wallis and Dunn’s multiple comparison test was performed. (F) Representative flow cytometry showing TIM-3 and LAG-3 expression in the different CAR populations on day 10 (left). Cumulative percentage of LAG3^+^/TIM3^+^ double positive cells (right). Paired data of 3-5 donors in 3-5 independent experiments with 1-2 internal replicates (different MOI transduction: 1.4 ± 0.6) are shown. Two-way ANOVA; Tukey’s multiple comparison test. <u>Abbreviations</u>. CRISPR, Clustered Regularly Interspaced Short Palindromic Repeats. HD, Hinge Domain. TMD, Trans-Membrane Domain. ICD, Intracellular Domain. LV, lentivirus. scFv, single chain variable fragment. CAR, Chimeric Antigen Receptor. bPD1-CAR, blocking anti-PD-1 CAR. nbPD1-CAR, non-blocking anti-PD-1 CAR. KO, knock out.

### In vivo expansion of anti-PD-1 CAR-T cells correlates with delayed HIV rebound

We next evaluated the ability of both anti-PD-1 CARs to control HIV replication in a hu-mice model where the leukocytes are susceptible to HIV infection (*21*–*23*). Immune reconstituted hu-mice were infected with the YU-2 HIV-1 strain from 16 weeks of age and 6-8 weeks later the infection was confirmed by RT-PCR. ART regimen was then initiated and once the mice suppressed, T cells were engineered either from the CD34^-^ fraction of the corresponding cord blood donors, and if the material was unavailable, from the spleen of autologous uninfected hu-mice (Fig. 5A). Considering that the success of CAR-T cell therapies relies on the fitness and synergic effect of CD4^+^ and CD8^+^ T cells (*24*, *25*) and that CD4^+^ T cells are susceptible to HIV infection, we first evaluated additional editing strategies conferring HIV resistance. Thus, we tested the influence of editing the CD4, CCR5 or CXCR4 receptors and/or a combination thereof on the susceptibility to HIV infection (Fig. S7A, B). To address this issue, edited cells were exposed to CCR5 or CXCR4 tropic pseudoviral strains expressing GFP or alternatively to HIV-BaL (CCR5-tropic lab-derived HIV variant) or HIV-IIIb (CXCR4-tropic lab-derived HIV variant) replication competent viruses (Fig S7C, D). In both cases, CD4 editing strongly reduced the susceptibility to HIV infection. Thus, we decided to target only the CD4 locus in addition to the PD-1 +/- TRBC for T cells expanded from the CD34^-^ fraction (to prevent xenogeneic reactions). Double (CD4-PD-1) and triple (CD4-PD-1-TRBC) editing efficiencies were variable ranging from 50-95% (Fig. S8).

**Fig. 5.**
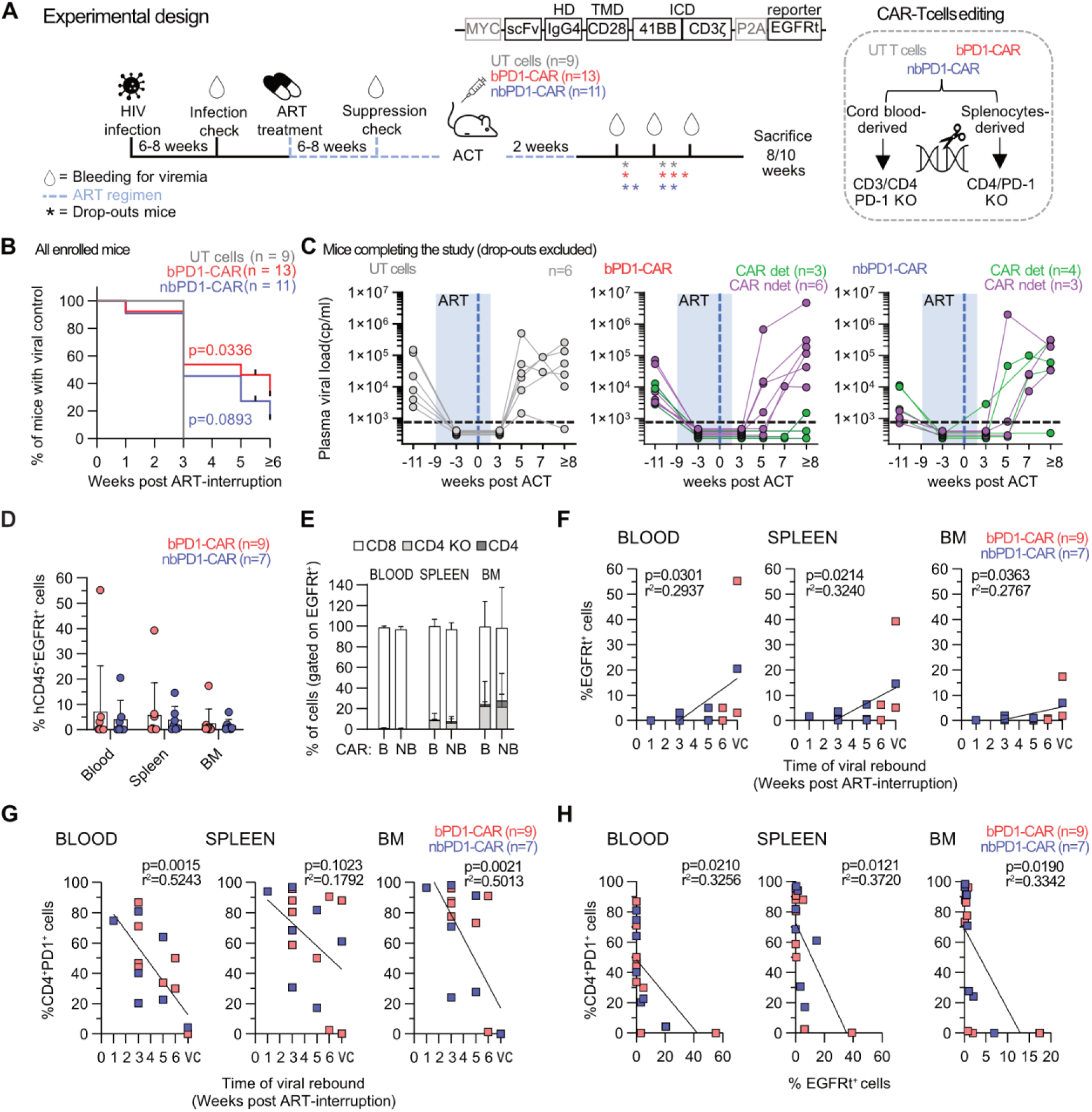
Anti-PD-1 CAR-T cells expansion *in vivo* correlates with depletion of CD4^+^PD-1^+^ cells and delayed viral rebound in humanized HIV infected mice. (A) Experimental design. Mice received adoptive cell transfer (ACT) of anti-PD-1 CARs versus untransduced (UT) T cells under ART, that was interrupted after two weeks (3 independent experiments). The reporter used for detecting CAR positive cells was EGFRt. (B) Kaplan-Meyer curve showing the percentage of mice maintaining viral control. All mice were included in the analysis. Log-rank (Mantel-Cox) statistical test was performed using the UT control condition as reference. (C) Plasma viral load overtime (UT, n=9; bPD1-CAR, n=9; nbPD1-CAR, n=7). (D) Percentage of CAR-T cells (defined as EGFRt^+^ gated in CD45^+^ cells) in the spleen, bone marrow and blood at the time of sacrifice. Mean ± SD is shown. bPD1-CAR, n=9, nbPD1-CAR, n=7. (E) Percentage of CD8^+^, CD4^+^ and CD4^-^ CAR-T cells detected in the spleen, bone marrow and blood (bPD1-CAR, n=3, nbPD1-CAR, n=4). Mean ± SD is shown. (F) Correlation between the percentage of CAR-T cells (defined as CD45^+^EGFRt^+^ cells) and the time of viral rebound (bPD1-CAR, n=9, nbPD1-CAR, n=7). Simple linear regression was used. VC means viral control. (G) Correlation between CD4^+^PD-1^+^ cells (gated in huCD45^+^EGFRt^-^ cells) and the time of viral rebound (bPD1-CAR, n=9, nbPD1-CAR, n=7). Simple linear regression was used. (H) Correlation between CD4^+^PD-1^+^ cells (gated in huCD45^+^EGFRt^-^ cells) and CAR-T cells detection (bPD1-CAR, n=9, nbPD1-CAR, n=7). Simple linear regression was used. <u>Abbreviations</u>. HIV, Human Immunodeficiency Virus. ART, anti-retroviral therapy. ACT, adoptive cell transfer. HD, Hinge Domain. TMD, Trans-Membrane Domain. ICD, Intracellular Domain. scFv, single chain variable fragment. CAR, Chimeric Antigen Receptor. bPD1-CAR, blocking anti-PD-1 CAR. nbPD1-CAR, non-blocking anti-PD-1 CAR. KO, knock out. UT, untransduced T cells. EGFRt, truncated Epidermal Growth Factor Receptor. VC, viral control.

ACT was performed under ART, that was interrupted after 2 weeks and viremia was measured by qPCR 1,3,5 weeks after ART interruption (Fig. 5A). Notably, viral rebound, defined by serum HIV level exceeding 500 HIV RNA copies/mL, was delayed in anti-PD-1 treated mice as compared to controls (p=0.0336 for the bPD1-CAR treated mice and p=0.0893 for the nbPD1-CAR) (Fig. 5B). Importantly, three mice (one nbPD1-CAR and two bPD1-CAR) showed undetectable viral load until week 6-8 after ART interruption (Fig. 5C). Interestingly, viral control was more efficient in the bPD1-CAR treated group and in mice with detectable CAR-T cells in organs. Notably, CAR-T cells were detectable in the spleen, bone marrow and/or blood of 33% of bPD1-CAR treated and 57% of the nbPD1-CAR treated mice up to 70 days after ACT, predominantly CD8^+^ CAR-T cells (Fig. 5D, E, Table S1-2). Importantly, we observed a significant positive correlation between viral control and CAR-T cell detection (Fig 5F) and negative correlations between PD-1 detection and viral control (Fig. 5G, Fig. S9A) or CAR-T cells detection (Fig 5H and Fig. S9B), suggesting that the control of HIV replication was indeed conferred by anti-PD-1 CAR-T cells.

### Anti-PD-1 CAR-T cells cause a disruption of the lymphoid architecture correlating with HIV RNA clearance

To better understand the mechanisms involved in the sustained control of HIV replication, we performed additional immunohistological analysis of the spleen of mice with sustained control of HIV replication (n=3), mice with documented viral rebound (n=4) and mice treated with untransduced T cells were used as controls (n=6). The spleens of control mice exhibited peculiar lymphoid structures characterized by close interactions of B cells, CD4^+^ T cells and CD8^+^ T cells (Fig. 6A and Fig. S10). Interestingly, mice with sustained control of HIV replication harbored a significant reduction of CD4^+^ PD-1^+^ T cells in the CD20-enriched zones, as well as a significant depletion of CD20^+^ B cells in CD4-enriched regions as quantified by histo-cytometry (Fig. 6B, and Fig. S11), which correlated with a significantly lower clustering of the B cells, indicating a disruption of the lymphoid structure (Fig. 6C). For one bPD1-CAR treated mouse, we found a strong proliferation of PD-1^+^ and Grzb^+^ CD8^+^ T cells (Fig. 6B and Fig. S12). To further confirm the control of HIV replication, we performed RNAscope studies on three animals, using one probe specific for the WPRE element of the CAR lentiviral vector and a second one for HIV RNA. We quantified both CAR-specific and HIV signals and confirmed the absence of cells harboring HIV RNA transcripts in the responder mouse S11, which is consistent with the blood qPCR results (Fig. 6D, E, Fig. S13). Importantly, in the S5 non-responder mouse, the HIV RNA^+^ cells were mainly CD4^+^ T cells, and the proportion of HIV RNA^+^ cells among the non-CD8 CAR^+^-T cells was less than 0.4% in both treated mice (S5 and S11). This suggests that CD4 editing was sufficient to protect CAR-T cells from HIV infection (Fig. S13). In conclusion. our *in vitro* and *in vivo* findings were further supported by in situ studies, collectively validating the efficacy of our CAR-T cells against HIV-infected cells.

**Fig. 6.**
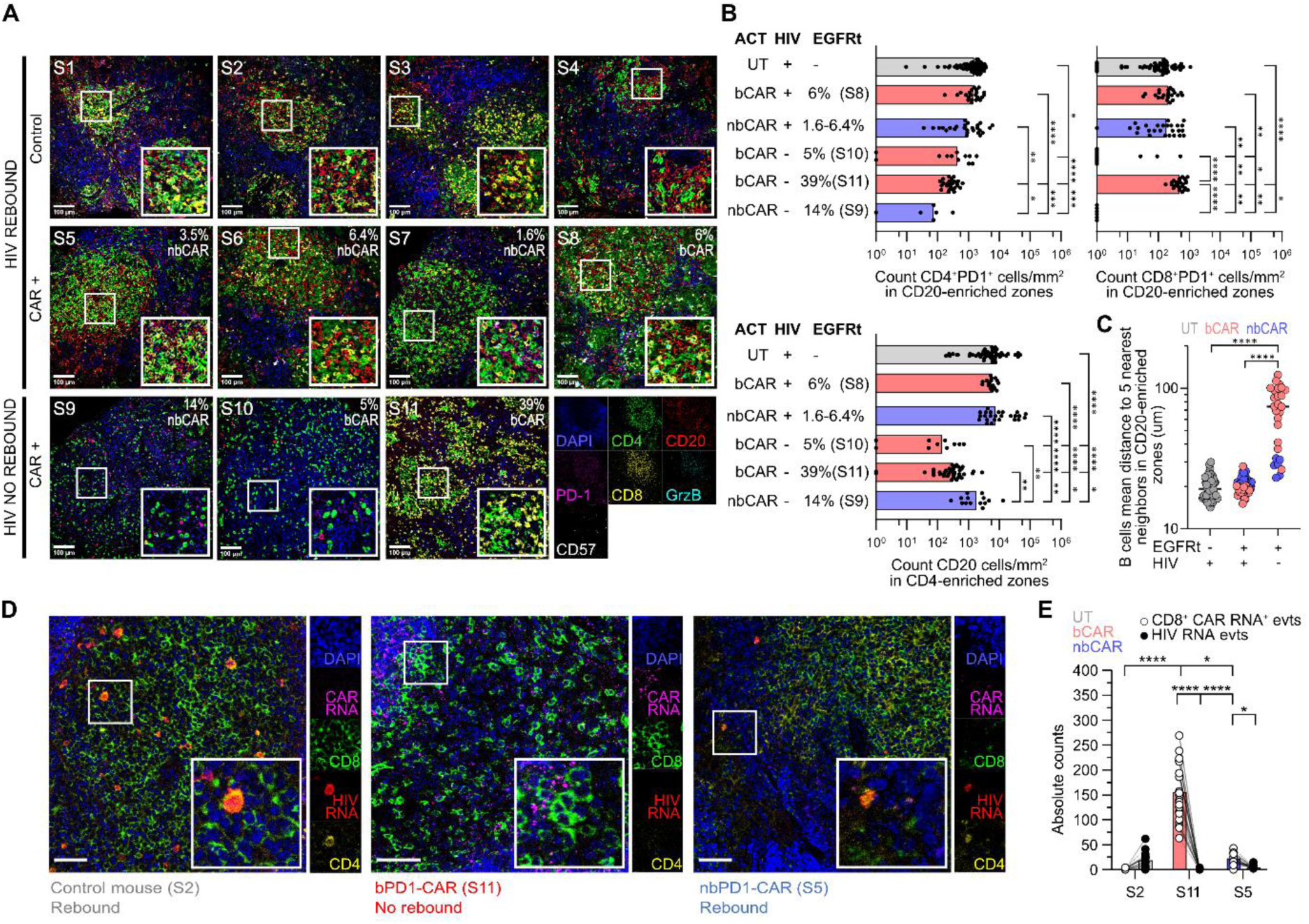
Anti-PD-1 CAR-T cells disrupts the lymphoid architecture in humanized mice spleen which correlates with HIV RNA tissue clearance. (A) Immunofluorescence multicolour confocal images of spleens of 4 control and 7 treated hu-mice with detectable CAR-T cells (untransduced treated n= 4, nbPD1-CAR-T treated n=4, bPD1-CAR-T treated n=3) 8 to 10 weeks after ACT. The following staining were performed: DAPI (blue), CD4 (green), CD20 (red), PD-1 (magenta), CD8 (yellow), GrzB (cyan) and CD57 (gray). Scale bar, 100 μm. (B) Histo-cytometry analysis showing CD4^+^PD-1^+^, and CD8^+^PD-1^+^ cell counts/mm^2^ in the CD20-enriched areas and CD20^+^ cell counts/mm^2^ in the CD4-enriched zones. Each dot represents a CD20 or CD4-enriched region (n=154 and n=158). Median is shown. Kruskal-Wallis and Dunn’s multiple comparison test. All control mice (n=6) and treated mice with detectable CARs (n=7) that survived until the end of the experiment were included. (C) Mean distance between each B cell and its 5 nearest neighbors in the CD20-enriched regions in the 13 spleens based on IF staining data. Median is shown. Kruskal-Wallis and Dunn’s multiple comparison test was performed. (D) Combined IF and RNAscope for CD8 (green), DAPI (blue), HIV RNA (red) and CAR RNA (magenta) performed on the spleen of one representative control mouse (S2), one HIV suppressed responder mouse (S11) and one non-responder mouse with viral rebound (S5) 8 weeks after ACT. Scale bar is 50 μm. (E) Histo-cytometry analysis showing absolute counts of CAR RNA events (gated on CD8^+^ cells) and HIV RNA events for each mouse tissue. Two-way ANOVA, Tukey’s multiple comparison test. <u>Abbreviations</u>. S, spleen. bCAR, blocking anti-PD-1 CAR. nbCAR, non-blocking anti-PD-1 CAR. ACT, Adoptive Cell Transfer. UT, untransduced T cells. EGFRt, truncated epidermal growth factor receptor.

## DISCUSSION

In this study, we demonstrate the feasibility of depleting subpopulations of human T cells expressing PD-1 in both ex-vivo cultures and hu-mice. Functionally, CAR-T cell persistence, particularly with the bPD1-CAR, correlated with HIV viral suppression, highlighting the potential of this strategy to reduce the HIV reservoir. Our in-situ analysis showed a disruption of the lymphoid architecture in the white pulp area of the spleen. This finding was consistent across the two anti-PD-1 CARs despite the lower sensitivity of the nbPD1-CAR. Finally, we showcase that the CAR epitope binding site modulates T-cell avidity, sensitivity, and trogocytosis capacities. Thus, adequate scFv selection when designing CARs could potentially reduce the burden of acute toxicities related to CAR-T cell therapies.

Our findings build upon prior evidence that PD-1^+^ CD4^+^ T cells serve as a key viral reservoir in lymphoid tissues (*14*). By engineering CAR-T cells to target PD-1^+^ cells, we propose an alternative approach to anti-HIV envelope-based CARs for eliminating HIV-infected cells, that remain limited by the constant mutations of the virus. This approach has also been evaluated by the group of Corey et al., who infused anti-PD-1 CAR-T cells in SIVmac239–infected rhesus macaques. They report a selective loss of CD4^+^ PD-1^+^ Tfh cells correlating with suppression of SIV in the germinal centers (*26*). Yet, the non-selective nature of the anti-PD-1 CARs led to the depletion of memory CD8^+^ T cells involved in the immune surveillance against HIV (*27*, *28*). Thus, this depletion induced paradoxically an immunosuppressive state and accelerated HIV rebound in the extra-follicular reservoir upon ART interruption (*26*). This limitation may be addressed by tuning down the CAR avidity or affinity as Tfh express substantially higher level of PD-1 or by co-injecting HIV-specific engineered T cells. Alternatively, the specificity could be improved by generating dual receptor either specific for the HIV reservoir such as CD32a (*29*, *30*), CXCR3 (*14*) or for other surface makers highly expressed on Tfh.

Interestingly, Eichholz et al. showed that only half of the treated Monkeys had persistent anti-PD-1 CAR-T cells (*26*), a similar percentage to our study. Despite an extensive analysis of the different confounding factors including the type of engineered T cells (CD34-cord blood cells versus splenocytes), the CD45 reconstitution level, the blocking/non-blocking nature of the CAR or the CD4 to CD8 ratios or the viral loads before ACT, we were not able to identify strong predictive biomarker (Table S3). It is therefore tempting to speculate that the level of PD-1 expression within the tissue, which may not correlate with the level of reconstitution, could influence the expansion and persistence of the CAR-T cells. HIV infection, ART, and CAR-T cell proliferation in lymphopenic environments could lead to different PD-1 expression level across mice (*31*). Importantly, suppressed HIV mice showed a selective expansion of CD8^+^ CAR-T cells, which is consistent with clinical studies demonstrating the importance of this population in predicting oncological responses (*25*, *32*). In future studies, we should evaluate the enrichment of CD7^+^CXCR3^+^ CAR-T cells before ACT as it has been recently shown to correlate with long-term CAR-T cell persistence and efficacy (*33*).

Herein, we show that our anti-PD-1 CAR-T cells can disrupt the lymphoid architecture in association with a depletion of CD4^+^ PD-1^+^ cells. Indeed, PD-1^high^ Tfh cells are essential to maintain the spatial organization of germinal centers and the follicles structure in secondary lymphoid organs (*34*, *35*). Interestingly, Bui et al., by infusing anti-CD20 CAR-T cells in SIV-infected monkeys, efficiently ablated B cell follicles and thereby Tfh cells. This depletion significantly reduced the splenic HIV reservoir, as did anti-PD-1 CAR-T cells (*36*). Furthermore, NK cells engineered with a fusion PD-L1 chimeric receptor depleted Tfh cells selectively, which inhibited B cell proliferation and antibody production (*37*). Altogether, these results underscore the important interplay between Tfh and B cells and the therapeutical potential of anti-PD-1 CAR-T cells to induce an immune reset of B cells follicles, a strategy which could be investigated in autoimmune diseases.

The present study has several limitations. First, our data showed that we were not specific to Tfh as we observed a deletion of CD8^+^ T cells. Moreover, infected hu-mice do not develop an HIV-specific CD8^+^ T cell response (*23*, *38*), preventing us from evaluating the respective contribution of each CAR in such deletion. Secondly, we have not evaluated the contribution and duration of the ART on the efficacy of the CAR-T cells, as we arbitrarily decided to maintain ART for additional two weeks post-ACT. Third, our study predominantly found an expansion of anti-PD-1 CAR CD8^+^ T cells *in vivo*. Therefore, we cannot exclude the possibility that the fitness of the anti-PD-1 CAR CD4^+^ T cells was impaired due to the gene-editing strategy, possibly reducing the efficacy of the therapy. Indeed, combining CD4^+^ and CD8^+^ CAR-T cells has been shown to have a synergistic anti-tumor effect *in vivo* (*39*, *40*). Thus, future studies should investigate the importance of co-injecting CD4^+^ CAR T cells, considering the susceptibility of CD4^+^ T cells to HIV infection. Potential alternative approaches for conferring HIV resistance include engineering peptides that inhibit the virus-cell fusion entry or deleting the CD4 receptor using safer methods, such as CRISPR-based editing (*41*). Finally, even if humanized mouse models provide valuable insights into HIV latency and potential eradication strategies (*42*), they are not suitable to detect potential CAR-T cell-associated toxicities.

In summary, we have developed two novel anti-PD-1 CAR-T cell products that, when combined with other strategies or enhanced through engineering improvements to increase specificity, could be a step forward in achieving a cure for HIV. Thus, our approach, which targets cellular rather than viral antigens, has the major advantage of avoiding the virus’s natural capacity to mutate gp120 and escape immune surveillance. Finally, our strategy could pave the way for innovative therapies for alternative indications such as autoimmune diseases considering the essential role of Tfh in the humoral response.

## MATERIALS AND METHODS

### Study design

The study aimed to develop novel anti-PD-1 CAR T cells and evaluate their impact on the HIV reservoir in humanized mice. The experimental design included editing of human T cells using CRISPR-Cas9 technology and lentiviral transduction. The functional activity of CAR T cells was evaluated in hu-mice susceptible to HIV infection. The dataset was supplemented with flow cytometry and histological analyses, derived from adoptive cell transfer experiments in hu-mice. Details on the number of biological replicates and the number of independent experiments are indicated in the figure legends and Data File S1. An independent experiment includes independent isolation, manufacturing, and expansion of the primary cells. For *in vivo* studies, the group assignment was performed arbitrarily based on cell availability, aiming to have all groups represented (untransduced, nbPD1-CAR, and bPD1-CAR) for each donor. For *in vivo* data analysis, mice with less than 10% huCD45^+^ cells in the spleen were excluded. For ex vivo flow cytometry data analysis of mice, CAR detection was arbitrarily considered positive if more than 1% of huCD45^+^EGFRt^+^ events were detected.

### Human blood products

Buffy coats of deidentified human peripheral blood from healthy donors was purchased from the Swiss Transfusion Center. Density gradient centrifugation with Ficoll-Paque (Cytiva, Uppsala, Sweden, Cat. #17144003) was used to isolate peripheral blood mononuclear cells (PBMCs).

### T cell isolation and flow cytometry

Human CD3^+^, CD4^+^ or CD8^+^ T cells were obtained using the EasySep (1) Human T Cell Enrichment Kit, (2) CD4^+^ T Cell Isolation Kit and (3) CD8+ T Cell Enrichment Kit (StemCell Technologies, Vancouver, Canada, Cat. #19051) respectively following the manufacturer’s instructions from PBMCs or CD34-cord blood cells. For T cells isolated from mice, splenocytes were stained with anti-CD45-FITC, anti-CD4-AF700 and anti-CD8-APC-Cy7 or PB anti-human antibodies, and CD4^+^ and CD8^+^ T cells isolated by FACS sorting on a BD FACS Aria II (BD Biosciences, Franklin Lakes, NJ, USA). For flow cytometry staining, BD LSR Fortessa Cell Analyzer (BD Biosciences, Franklin Lakes, NJ, USA) was used for sample analysis and FlowJo v10.9.0 software (BD Biosciences, Franklin Lakes, NJ, USA) for data processing. All antibodies used in this study are listed in Table S4.

### Lentivirus production

Third generation lentiviruses were produced in HEK293T cells using 11.25 µg of the transgene containing vector, 2.8 µg of packaging pRSV-Rev Addgene Plasmid (Cat.#12253) and 7.3 µg of packaging pMDLg/pRRE Addgene Plasmid (Cat.#12251) and 3.95 µg of envelope expressing pMD2.G Addgene Plasmid (Cat.#12259) for transfection. Third generation lentiviral backbone was kindly gifted by Kirsten Scholten (Center of Experimental Therapy, Department of Oncology, University Hospital of Lausanne (CHUV), Switzerland). 16h after transfection cells were washed, 24 and 48 hours later supernatant was collected and filtered with a 0.45µm filter. Supernatant was then concentrated by ultracentrifugation at 68000g for 2hours at 12°C and stored at -80°C. The CCR5-tropic and CXCR4-tropic HIV-derived vector encoding for EGFP (HIVR5GFP and HIVX4GFP) were obtained as a kind gift from Nicolas Manel (Institut Curie, Paris, France) and amplified as previously described (*43*, *44*).

### Human primary T cells transduction and expansion

Human primary T cells were activated with anti-human CD3/CD28 Dynabeads (Gibco, Waltham, MA, USA, Cat. #11131D) at 1:2 bead:T cell ratio in X-Vivo 15 medium (Lonza, Basel, Switzerland, Cat. #02-053Q) with 5% human type AB serum from male donor (Pan-Biotech, Aidenbach, Germany, Cat. #P30-2901), 1% penicillin-streptomycin (BioConcept, Allschwill, Switzerland, Cat. #4-01F00-H), 55 μM 2-mercaptoethanol (Gibco, Grand Island, NY, USA, Cat. #21985-023), and 10 mM N-acetyl-L-cysteine (Sigma-Aldrich, Saint-Louis, MO, USA, Cat. #A9165-25G). After 18-20 hours from activation, lentiviral transduction was performed by adding the lentivirus of interest in the T cell culture at MOI=1.4 ± 0.6. After 24 hours from lentiviral transduction cells were washed and cultured for the rest of the expansion in Roswell Park Memorial Institute (RPMI) medium (Gibco, Waltham, MA, USA, Cat. #61870-010) supplemented with 10% FBS (Sigma-Aldrich, Saint-Louis, MO, USA, cat. #F7524), 1% non-essential amino acids (NEAA, Gibco, Grand Island, NY, USA, cat. #11140-050), 10 mM hepes buffer solution (Gibco, Paisley, UK, cat. #15630-056), 1 mM sodium pyruvate (Gibco, Waltham, MA, USA, cat. #11360-039), and 1% penicillin-streptomycin. Medium was supplemented with 30 IU/mL recombinant human IL-2 (rhIL-2, Miltenyi Biotec, Bergisch-Gladbach, Germany, Cat. #130-097-746) for T cells or CD4^+^ T cells, 100 IU/mL for CD8^+^ T cells when expanded alone. For the *in vivo* experiments, the T cell medium was enriched with 5 ng/mL recombinant human IL-7 (rhIL-7, Miltenyi Biotec, Bergisch-Gladbach, Germany, Cat. # 130-095-362), and 5 ng/mL recombinant human IL-15 (rhIL-15, Miltenyi Biotec, Bergisch-Gladbach, Germany, Cat. #130-095-764) and 30IU/ml IL-2. Primary T cells were cultured at a confluency of 1-2×10^6^ cells/mL at 37°C with 5% CO_2_ and cytokines renewed every other day. On day 7 of expansion anti-CD3/CD28 beads were removed, and T cells were rested overnight in complete medium before performing functional assays. For the *in vivo* experiments cytokines were maintained for all duration of expansion and washed out before ACT.

### Humanized mice studies

All animal experiments were reviewed and approved by the Cantonal Veterinary Office and performed in accordance with local guidelines and Swiss animal protection law. Use of human fetal liver tissue was approved by the cantonal ethical committee of Zurich, Switzerland. Hu mice were generated as previously described (*45*). Briefly, Newborn NOD.Cg-Prkdc^scid^ Il2rg^tm1Wjl^/SzJ (NSG) mice were irradiated with 1 Gy 1-3 days after birth and subsequently transplanted with 1.0 ± 0.5 x10^5^ CD34^+^ cord blood cells via intrahepatic injection. Immune reconstitution in female and male was evaluated at 16 weeks of age by staining the peripheral blood for huCD45.

Hu-mice with human engraftment level of CD45^+^ cells > 5% were used for further experiments. For HIV infected mice, an intraperitoneal injection at a TCID50 of 2×10^5^ with HIV-1 YU-2 was done as previously reported (*38*). Briefly, viral load was measured in the blood 6 weeks after infection using a specific qPCR assay against *vif* found in HIV-1 clade B strains. RNA was isolated from 20ul of plasma and eluted in 60ul AVE buffer using the QIAamp Viral RNA Mini Kit (Qiagen, Hilden, Germany Cat. #52904). Retrotrascription and pre-amplification were performed in one step using the SuperScript™ III Platinum™ One-Step qRT-PCR Kit w/ROX (Thermo Fisher Scientific, Waltham, MA, USA, Cat. #11745100). Primer for *vif* target region are: GGTCTGCATACAGGAGAAAGAG and GCTAGTTCAGGGTCTACTTGTG. PCR product was then diluted 1:10 in RNAse free water and used for real-time qPCR with the same primers and the following probe: /56- FAM/ACTGGCATT/ZEN/TGGGTCAGGGAGTC/3IABkFQ/. For real-time qPCR we used FastStart Essential DNA Probes Master (Roche, Roche Holding AG, Indiana, USA Cat.# 6402682001). Standards were generated by using an HIV-1 culture supernatant with a known copy number. Once infected the mice received a combination of raltegravir, tenofovir and emtricitabine as cART for viral load suppression.

### Mouse tissues preparation for flow cytometry and ex vivo analysis

For ex vivo analysis mice blood and spleen mononuclear cells were isolated by density gradient centrifugation using Lymphoprep™ (StemCell Technologies, Vancouver, Canada, Cat. ##18060), after smashing the organ on 70 µm cell strainers. For bone marrow processing, bones were cut and flushed with a syringe, cell suspension passed through 70 µm cell strainers and red blood cells removed using ACK lysing buffer (Lonza, Basel, Switzerland, Cat. #10-548E). Cells were washed twice in MACS buffer (PBS with 2 mM EDTA and 2% FBS) and stained with anti-human CD45 FITC, CD3 BV786, CD4 PE-Cy7, CD8 PB, EGFR PE, PD-1 APC, Live/Dead APC-Cy7. Samples were fixed with 1% PFA for 40 minutes at 4°C before analysis at the BD LSR Fortessa Cell Analyzer (BD Biosciences, Franklin Lakes, NJ, USA).

### Statistical analysis

For most in vitro experiments with human primary cells, each replicate was a unique healthy donor. Depending on material availability and cells count numbers, cells from one donor were transduced with different MOIs and were considered as independent replicates. For *in vivo* experiments, each replicate was a single mouse. GraphPad Prism 10 software was used for all statistical analysis. Data distribution was assessed using the Shapiro-Wilk test. For normally distributed data, statistical differences between two groups were determined using two-tailed parametric Student’s t tests, whereas comparisons among three or more groups were conducted using one-way or two-way analysis of variance (ANOVA) with Tukey’s post hoc correction when comparing all groups between them or Dunnett’s post-hoc correction when comparing all groups to control group. For data that did not pass the normality test, statistical differences between three or more groups were performed using Kruskal-Wallis tests with Dunn’s post hoc analysis. Correlations were analysed with a simple linear regression model. Kaplan-Meyer survival curves for viral load were analyzed with Log-rank (Mantel-Cox) statistical test.

Specific test and statistical method are described for each analysis in the figure legend. Statistical significance values of the stars are the following: * P ≤ 0.05, ** P ≤ 0.01, *** P ≤ 0.001, **** P ≤ 0.0001. Only statistical differences are reported. No outliers were excluded.

## Acknowledgments

We thank the Center for Immunotherapy and Vaccinology, Division of Immunology and Allergy at the University Hospital of Lausanne (CHUV), for their precious support.

## Author contributions

Conceptualization: Y.D.M., G.P.

Supervision: Y.D.M.

Designed experiments: L.E, Y.D.M.

Performed experiments: L.E., R.B., N.K., S.G., M.O., F.P., A.A.S., O.A.A., R.C., R.P.O.

Analysed data: L.E., Y.D.M., DC.C., S.G., M.O., C.B., C.P., M.F.

Provided reagents and advice: C.P., M.P., R.B., F.P., C.F., L.P., R.S., N.K., S.G., K., R.C., E.L., O.Y.C. G.P

Wrote the original draft: L.E., Y.D.M.

Reviewed and edited the manuscript: all

All authors approved the manuscript.

## Data and materials availability

All data associated with this study are present in the paper or the Supplementary Materials. All reasonable requests for data and code will be fulfilled.

## List of Supplementary Materials

Materials and Methods

Fig. S1 to S13

Table S1 to S6

References (*46-59*)

