## Supplementary materials for "Targeting PD-1^+^ T-cells with Chimeric Antigen Receptors to reduce the HIV Reservoir"

Laura Ermellino *et al.*

**This file includes:**

Materials and Methods
Figs. S1 to S13
Tables S1 to S6
References (*46-59*)

**Supplementary materials and methods**

**Cell lines generation**

The PD-1 gene in Jurkat and K562 cell lines was deleted using guides targeting the exon 2 of PDCD1. The crRNA sequence was kindly shared by Dr. Marcela Maus, Massachusetts General Hospital, Harvard University, Boston, MA (*46*). Single cells were plated, and clones were screened for indel insertions in the PDCD1 locus. Successful editing was confirmed by sanger sequencing and using the ICE CRISPR Analysis Tool (Synthego, Redwood City, CA, USA) (*47*). One PD-1 KO clone for each cell line was selected to produce all other cell lines reported in the result section. To obtain differential PD-1 expression, the Jurkat PD-1 KO clone was transduced with a lentivirus encoding for PD-1 under a CD4 promoter. Single cell clones were plated to obtain stable PD-1 expression ranging from 424 to 6047 molecules. PD-1 molecules were counted using BD Quantibrite™ Beads PE Fluorescence Quantitation Kit (Cat. #340495). The design of alternative lentiviral constructs is described in the results section, and all are under an EF-1 promoter. Cell lines were routinely tested and confirmed negative for mycoplasma.

**Antibodies prediction model**

Model of the Fab fragments of antibodies A35795 (A35) and 135c139d6 (135c) were generated using AlphaFold (*48*, *49*). For graphical representation, models and crystal structure of hPD-1/hPD-L1 complex (PDB ID: 4ZQK) and 135c/hPD-1 (PDB ID:6HIG) were aligned and Fabs model were placed. For A35 positioning, we used HADDOCK server and ClusPro-AbEMap (*50*, *51*). Chimera X was used for visualization and image generation.

**Antibody production and binding assay**

Antibodies were produced transfecting ExpiCHO cells with 1:1 ratio of plasmid AbVec2.0-IGHG1 (Addgene Plasmid Cat.#80795) containing the heavy chain sequence and the plasmid AbVec1.1-IGKC (Addgene Plasmid Cat.#80796) containing the light chain sequence at the Protein Production and Structure Core Facility of the Swiss Institute of Technology in Lausanne

(EPFL, Switzerland). Fabs were produced by digesting the IgG with papain using the Pierce™ Fab Preparation Kit (Thermo Fisher Scientific, Waltham, MA, USA, Cat. # 44985). scFv was cloned and produced as previously described (*52*).

PD-1$^{hi}$ transgenic Jurkat cells were used to evaluate antibodies (full, Fab and scFv) binding affinities. After 30 minutes at 4°C, cells were washed and stained with Goat anti-Human IgG (H+L) Cross-Adsorbed Secondary Antibody, Alexa Fluor™ 568 for IgG and Fab detection (Thermo Fisher Scientific, Waltham, MA, USA, Cat. # A-21090) and a PE anti-His Tag Antibody for scFv detection (BioLegend, San Diego, USA Cat.# 362603) respectively. For the competitive assays, cells were incubated with the Nivolumab or Pembrolizumab (0.3 to 10 µg/ml) for 30 min at 4°C followed by an 30 min incubation the biotinylated anti–PD-1 Abs (clone A35 or 135C) at 2 µg/ml and 30 min incubation with BD Pharmingen™ PE Streptavidin (BD Biosciences, Franklin Lakes, NJ, USA Cat.#554061). Clinical lots of Pembrolizumab (Keytruda, Merck) and Nivolumab (Opdivo, Bristol-Myers Squibb) were obtained through the Hospitalier Universitaire Vaudois.

**Biolayer interferometry**

BLI experiments were performed in PBS at 30 °C using an Octet K2 instrument (ForteBio) as previously described (*53*). Kinetic analysis of A35/135C IgG, Fab and scFv binding to PD-1 dimer protein was performed by biolayer interferometry immobilizing 3 µg/ml of PD-1 biotinylated dimer on streptavidin biosensors (Sartorius, Göttingen, Germany, Cat.18-5019#) dipped into a solution of the Fab/scFv at different concentrations diluted in PBS (ranging from 40 to 240 nM), and the nm shift was recorded on the Octet. Analysis was performed using the Octet software with 1.1 analyte fitting for the interaction with Fabs and scFvs.

**RNP formulation and electroporation protocols**

Cas9 protein was synthetized by the Protein Production and Structure Core Facility of EPFL (Switzerland). CRISPR-Cas9 guides were reconstituted as previously described (*54*, *55*). In brief, lyophilized crRNAs were resuspended in IDT nuclease-free duplex buffer at a 160 µM final concentration and stored at -80°C. Guides were reconstituted by mixing crRNA and tracrRNA at 1:1 ratio and incubated at 37°C for 30 minutes for annealing. Then, 45µM recombinant Cas9 was added to the 80 µM RNA at a 1:1 ratio and incubated at 37°C for 15 minutes. RNPs were kept at 4°C up to two weeks. Table S5 provides a list of the sequences of the guides used in this study. Human primary T cells were electroporated on day 0 before activation using the program EH-115 in the Lonza 4D 96-well electroporation system. After electroporation, cells were recovered with pre-warmed complete medium and rested for 10 minutes at 37°C.

**In vitro functional assays**

Functional assays were performed on day 8 of expansion after overnight resting without cytokines. For luciferase-based killing assay, $0.02 \times 10^6$ Jurkat cells were co-cultured with the CAR-T cells at different E:T ratios. After 24h luciferase assay reading was performed as previously described (*56*). Briefly, cells were lysed for 20 minutes by shaking at 300 rpm with 50 µL of harvesting buffer (50 mM 2-morpholinoethanesulfonic acid sodium (NaMES, pH 7.8, Sigma-Aldrich, Saint-Louis, MO, USA, cat. #1061970100), 50 mM Tris-HCl (pH 7.8), 1 mM dithiothreitol (Thermo Fisher Scientific, Waltham, MA, USA, cat. #R0861) + 0.2% Triton X-100 (Thermo Fisher Scientific, Waltham, MA, USA, Cat. #9002-93-1) in distilled water). Then, 50 µL of luciferase assay buffer (125 mM MES, 125 Tis-HCl (pH 7.8), 25 mM magnesium acetate tetrahydrate (Sigma-Aldrich, Saint-Louis, MO, USA, cat. #M0631), 2.5 mM adenosine triphosphate (ATP, Thermo Fisher Scientific, Waltham, MA, USA, cat. #R0441) in distilled water) were added for one minute. Thereafter 50 µL of luciferin buffer (1 mM D-luciferin (Thermo Fisher Scientific, Waltham, USA, cat. #88292) in 5 mM potassium dihydrogen phosphate (Merck, Darmstadt, Germany, cat. #1.04873)) were added to the lysate. Luminescence was measured on a Synergy H1 Hybrid reader (BioTek, Winooski, VT, USA).

For the activation assay, $0.1x10^6$ CAR-T cells were co-cultured with 120-Gy-irradiated K562 cell line at 1:1 E:T ratio. CD25 and CD71 was assessed 48h later by flow cytometry.

**HIV virus**

Human Immunodeficiency Virus Type 1 (HIV-1) BaL and IIIb strains were obtained from the National Institutes of Health (NIH) HIV Reagent Program. HIV-YU2 strain for hu-mice infection was purchased from the Centre for AIDS Reagents (National Institute for Biological Standards and Control, NIBSC, UK Cat.# 100 840).

**In vitro HIV-1 infection assays**

Primary $CD4^+$ T cells were isolated and edited for CD4/CCR5/CXCR4 in all different combinations of single, double or triple KO. After 7 days of expansion, edited T cells were re-stimulated with anti-CD3 mAb plate-bound and 3 days later infected with replication incompetent (R5 or X4 tropic) GFP+ HIV. Infection rate was evaluated by flow cytometry four days later. For the virological assay, edited T cells were sorted on day eight before re-stimulation. Cells were then infected on day 11 with replication competent HIV strains, HIV-BaL (R5 tropic) or HIV-1 IIIB Strain (X4 tropic). After 7 days in culture HIV RNA was quantified in the supernatant.

**Mouse tissue preparation and staining for immunofluorescence**

Half of the mice spleen was harvested for histological analyses. Tissues were fixed in 10% formalin overnight and then stored in 1%formalin-PBS. 3-µm sections from paraffin embedded (FFPE) blocks were prepared. Deparaffinization, antigen retrieval and fluorescent staining procedures were performed on the Ventana Discovery Ultra Autostainer (Roche Diagnostics, Basel, Switzerland) as previously described (*57*). Briefly, staining procedure consisted of subsequent cycles of antibody incubations and blocking steps using the Opal blocking/antibody diluent solutions (Akoya Biosciences, Marlborough, MA, USA, cat. #ARD1001EA). Primary antibody incubation was 32 min followed by a 16 min incubation with secondary HRP-labeled antibodies and a final one with optimized fluorescent Opal tyramide signal amplification (TSA) dyes (Opal 7-color Automation IHC kit and Opal650 reagent pack, Akoya Biosciences, cat. #NEL821001KT and #FP1496001KT, respectively). Repeated antibody denaturation cycles were performed. Tissue sections were washed and stained with Spectral DAPI from Akoya Biosciences for 4 min, rinsed in water with soap and mounted using DAKO mounting medium (Agilent, Santa Clara, CA, USA, cat. #S302380-2). Details on the antibodies, clones and dilutions are listed in Table S6.

**Quantitative Imaging Analysis (Histo-Cytometry)**

Confocal images were quantitatively analysed through histo-cytometry procedure as previously described (*58*, *59*), using Imaris software version 9.9.0 (Bitplane). Shortly, 3-dimensional segmented surfaces (based on the nuclear signal) were generated with the Surface Creation module of Imaris. Quantitative data generated this way, containing values like average intensities, together with volume and sphericity of the surfaces were exported in Microsoft Excel format. Files were then converted to comma separated value (.CVS) files and imported into FlowJo (version 10) for further analysis. For quantification of the different cell subsets manual gating of CD20-enriched or CD4-enriched regions was performed based on cell density.

**Quantitative spatial analysis of B cells organization**

To assess the spatial distribution of B cells within tissue sections, we performed a nearest-neighbor distance analysis using single-cell spatial coordinates. The (X, Y) spatial coordinates of $CD20^{+}$B cells were extracted for each enriched region, with at least 20 evts, and exported into separate Excel files, with each file corresponding to a specific region. Multiple regions were analyzed from each donor. Cells annotated with “KM_label = 1” indicated the $CD20^{+}$ B-cell population and were retained for further downstream analysis. The data were then imported and processed into R (version 4.4.2) using the “readxl” and “dplyr” packages. The spatial coordinates were transformed into a planar point pattern object using the ppp() function from the “spatstat” package, with tissue boundaries defined by the coordinate ranges specified through owin(). To evaluate the local spatial structure, we calculated the Euclidean distance from each $CD20^+$ B cell to its five nearest neighbors (k = 5) using the nndist() function. The analysis returned a set of distance values corresponding to the five nearest neighbors for each cell. To quantify the overall spatial organization within each donor, we computed the mean of the nearest-neighbor distance values across all regions analyzed per donor.

**RNAscope**

RNAscope *in situ* hybridization for the CAR RNA and HIV RNA visualization was performed according to the manufacturer’s instructions using the RNAscope™ Multiplex Fluorescent Detection Kit v2 (Advanced Cell Diagnostics, ACD, Newark, CA, USA Cat.# 323110). After deparaffinization on the Ventana Discovery Ultra Autostainer (Roche Diagnostics, Basel, Switzerland) as described above, the sections were treated with RNAscope Hydrogen Peroxide for 10min at RT, followed by an antigen retrieval step at 100°C for 15min. Subsequently, sections were incubated with Protease III for 15min at 40°C in a HybEz hybridization oven (ACD) to permeabilize the tissue. To detect CAR-T cells, we used the RNAscope™ Probe-WPRE-O4-C2 (Advanced Cell Diagnostics, ACD, Newark, CA, USA Cat.# 540341-C2) recognizing the Woodchuck Hepatitis Virus Posttranscriptional Regulatory Element specifically present in the 3’UTR of the CAR lentivector. For HIV RNA detection we used RNAscope® Probe - V-HIV1-CladeB-O1-C1 (Advanced Cell Diagnostics, ACD, Newark, CA, USA Cat. #1120101-C1). To be able to visualize the RNA signals, we used the tyramide based detection system by Akoya (Opal 7-color Automation IHC kit, Akoya Biosciences, Cat. #NEL821001KT). Afterwards, slides were incubated in antibody blocking for 30 minutes and then the protein staining was performed. Applied antibodies were verified for their compatibility with RNAScope protocol. Samples were incubated for 90 minutes, RT, with conjugated anti-CD4 AF700 and anti-CD8 A647. The samples were then counterstained with DAPI, and mounting was performed thereafter as described above.

**Image acquisition**

Acquisition of the images was performed on a Leica Stellaris 8 SP8 confocal system running the LAS-X software, at 512 x 512 pixel density for overview of the full tissue and 1024-1024 for high resolution image using the 0.75x optical zoom and a 20X objective (40X objective for RNAscope). Leica LAS-AF Channel Dye Separation module (Leica Microsystems) was used to create and apply a compensation matrix. For RNAscope histocytometry analysis the whole tissue was divided into 18-25 squares for a systematic analysis of the number of $CAR^+$ RNA and $HIV^+$ RNA cells across the tissue

### Supplementary figures

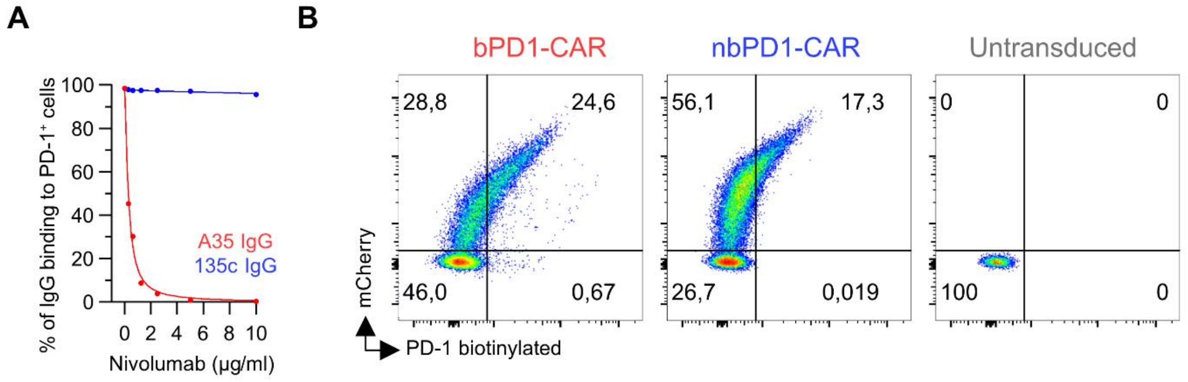

**Fig. S1. Competition binding assay and PD-1 binding to the CAR**

(A) Competitive binding assay with Nivolumab. Symbols are means of two independent experiments.

(B) Representative flow cytometry showing the binding of a PD-1 biotinylated protein (0.3µg/ml) to the anti-PD-1 CAR or untransduced T cells.

Abbreviations. IgG, Immunoglobulin G. PD-1, Programmed cell death protein 1.

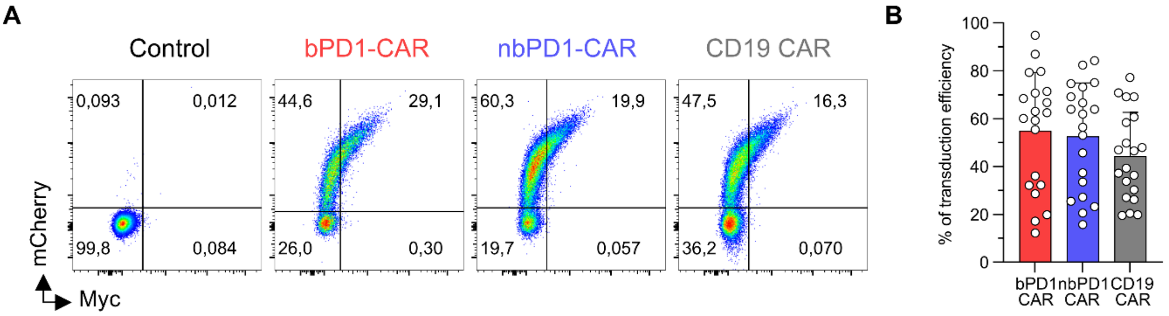

**Fig. S2. CAR expression and transduction level**

(A) Representative flow cytometry data showing mCherry and Myc co-expression on CAR-T cells.

(B) Cumulative data of transduction efficiency for donors reported in Figure 2D-E. The Mean ± SD values 8 donors, 8 independent experiments, 2-3 replicates transduced with different MOI is shown. Two-way ANOVA, Tukey's multiple comparison test.

Abbreviations. CAR, Chimeric Antigen Receptor. bPD1-CAR, blocking anti-PD-1 CAR. nbPD1-CAR, non-blocking anti-PD-1 CAR.

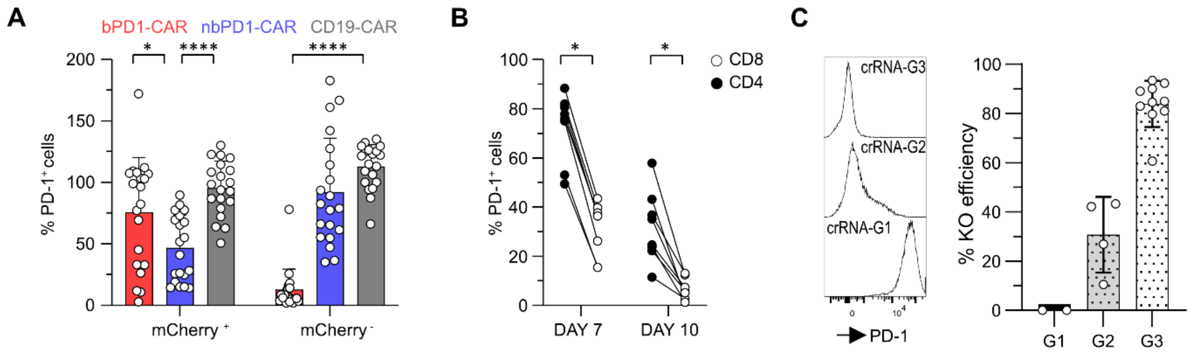

**Fig. S3. PD-1 expression in CAR-T cells and PD-1 knock-out efficacy**

(A) PD-1 expression in Cherry positive versus Cherry negative populations; Mean ± SD values of 8 donors, 8 independent experiments, 2-3 internal replicates transduced with different MOI is shown. Two-way ANOVA, Tukey's multiple comparison test.

(B) PD-1 expression in polyclonal $CD4^{+}$ and $CD8^{+}$ unedited T cells on day 7 and 10 of expansion. Mean of 8 donors, 8 independent experiments is shown. Multiple Paired T test.

(C) PD-1 editing tests with three different crRNAs and their efficiency (Mean ± SD values of n=2-10).

Abbreviations. PD-1, Programmed cell death protein 1. CAR, Chimeric Antigen Receptor. bPD1-CAR, blocking anti-PD-1 CAR. nbPD1-CAR, non-blocking anti-PD-1 CAR. KO, knock out. crRNA, CRISPR RNA.

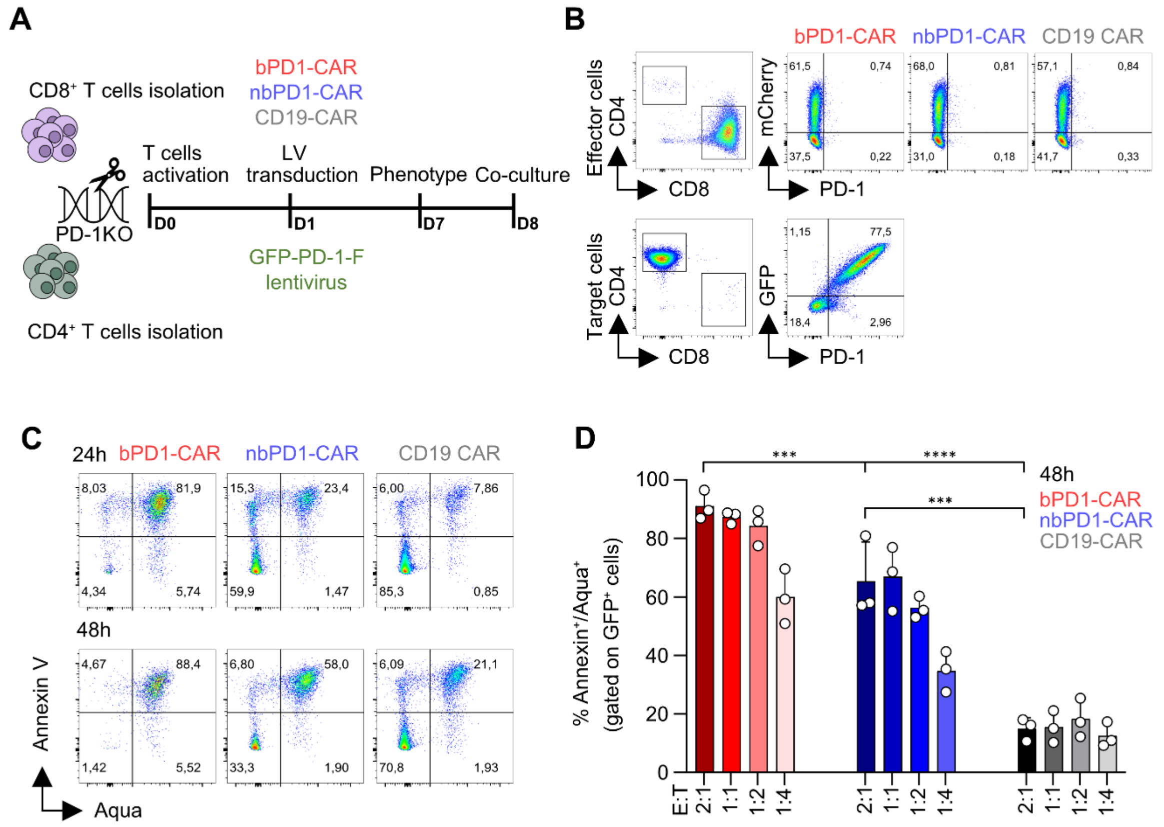

**Fig. S4. In vitro killing assay**

(A) Experimental design. CD8$^{+}$ and CD4$^{+}$ T cells were isolated, edited for PD-1, activated and expanded separately. CD8$^{+}$ T cells were transduced with different CAR constructs, while CD4 cells were transduced to stably express a PD-1-GFP-F fusion protein. Killing assay was performed on day 8.

(B) Representative flow cytometry of the phenotype of the target and effector cells used in the assay.

(C) Representative flow cytometry showing Annexin/Aqua staining in CD4$^{+}$GFP$^{+}$ target cells after 24h and 48h co-culture at 1:1 ration with CARs.

(D) Cumulative percentage of Annexin/Aqua$^{+}$ CD4$^{+}$PD-1$^{+}$ target cells in a FACS-based killing assay after 48h. Mean ± SD of 3 donors and 3 independent experiments is shown. Two-way ANOVA, Tukey's multiple comparison test.

<u>Abbreviations</u>. PD-1, Programmed cell death protein 1. CAR, Chimeric Antigen Receptor. bPD1-CAR, blocking anti-PD-1 CAR. nbPD1-CAR, non-blocking anti-PD-1 CAR. LV, lentivirus. KO, knock out.

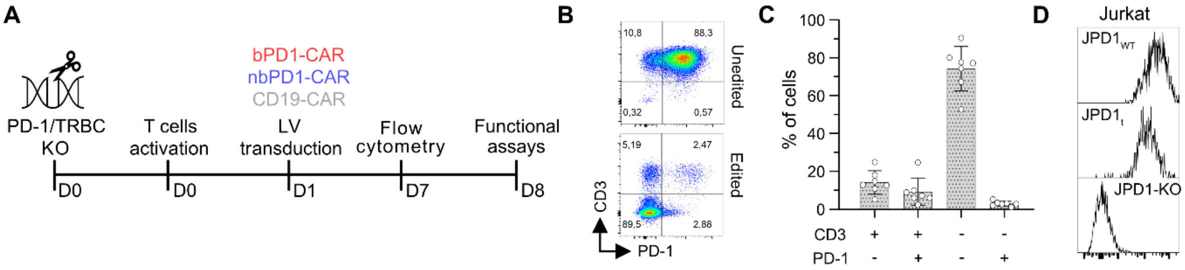

**Fig. S5. Experimental timeline and editing efficiency for functional assays**

(A) Experimental timeline of CAR T cells generation for functional assays.

(B) Representative flow cytometry staining showing CD3 and PD-1 on day 7.

(C) Cumulative data of editing efficiencies showing CD3 and PD-1. The Mean ± SD values of 7 donors is shown.

(D) PD-1 expression in the Jurkat cell lines used for luciferase-killing assay.

Abbreviations. CAR, Chimeric Antigen Receptor. bPD1-CAR, blocking anti-PD-1 CAR. nbPD1-CAR, non-blocking anti-PD-1 CAR. LV, lentivirus. TRBC, TCR beta chain. KO, knock out. JPD-1, Jurkat-PD-1. JPD-1-t, Jurkat-PD-1 truncated. UE, unedited.

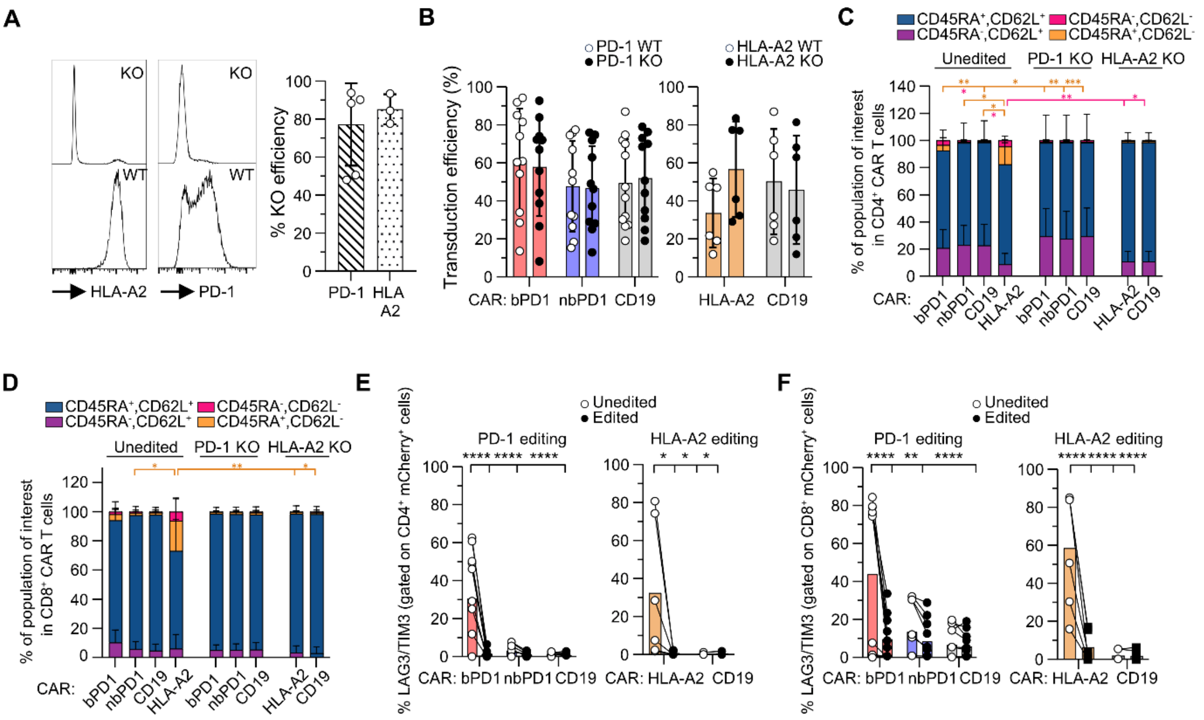

**Fig. S6. PD-1 and HLA-A2 editing and CAR-T cells differentiation and exhaustion profile**
(A) Representative flow cytometry staining showing PD-1 and HLA-A2 expression +/- editing (left). Cumulative data (right). For PD-1 KO, mean ± SD values of the untransduced condition of 5 donors in 5 independent experiments is shown. For the HLA-A2 KO mean ± SD values of the untransduced condition of 3 donors in 3 independent experiments is shown.
(B) Cumulative data showing the transduction efficiency +/- editing. Mean ± SD values of 3-5 donors in 3-5 independent experiments with 1-2 internal replicates (different MOI transduction: 1.4 ± 0.6) is shown. Two-way ANOVA, Tukey’s multiple comparison test.
(C-D) Cumulative data showing CD62L and CD45RA expression in edited versus unedited CD4$^+$ (C) and CD8$^+$ (D) CAR-T cells. The mean ± SD values of 3-5 donors in 3-5 independent experiments with 1-2 internal replicates (different MOI transduction: 1.4 ± 0.6) is shown. Kruskal-Wallis and Dunn's multiple comparison test was performed
(E-F) Percentage of LAG3$^+$TIM3$^+$ double positive CD4$^+$ (E) and CD8$^+$ (F) T cells in unedited versus edited CAR-T cells. Paired values corresponding 3-5 donors in 3-5 independent experiments with 1-2 internal replicates (different MOI transduction: 1.4 ± 0.6) are shown. Two-way ANOVA, Tukey’s multiple comparison test.
Abbreviations. CAR, Chimeric Antigen Receptor. bPD1-CAR, blocking anti-PD-1 CAR. nbPD1-CAR, non-blocking anti-PD-1 CAR. KO, knock out.

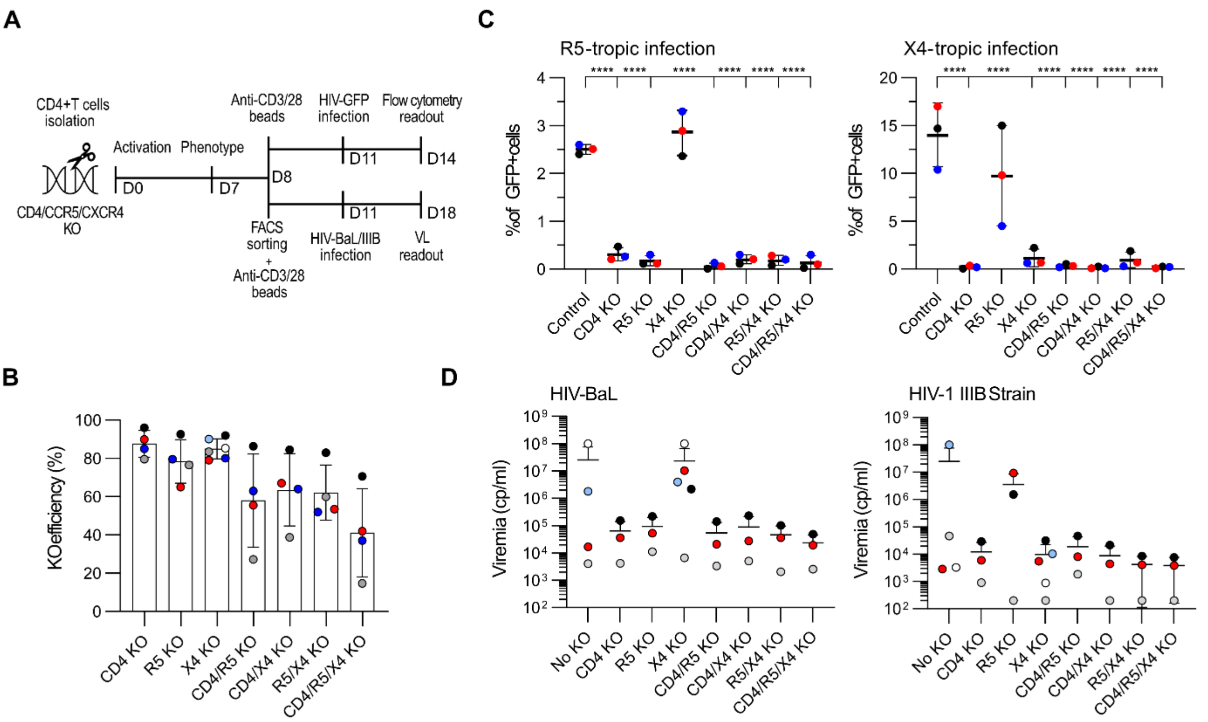

**Fig. S7. Generation of HIV resistant CD4⁺ T cells**

(A) Experimental timeline.

(B) KO efficiency for CD4, CCR5 and CXCR4 receptors in different combinations in CD4 primary T cells. Mean ± SD of 4-6 donors in 5 independent experiments.

(C) Percentage of GFP⁺ infected cells from a R5 tropic strain on the left and a X4 tropic strain on the right for each edited population. Mean ± SD values of 3 donors in 3 independent experiments. Comparison of each edited population with the unedited control is shown. Ordinary one-way ANOVA, Dunnett's multiple comparisons test was performed.

(D) Viral load (cp/ml) expressed in log scale measured in supernatant of CD4⁺ edited T cells 7 days after infection with HIV-BaL strain on the left or HIV-1 IIIB strain of the right. Mean ± SD values of 3-5 donors in 4 independent experiments. Comparison of each edited population with the unedited control is shown. Ordinary one-way ANOVA, Dunnett's multiple comparisons test was performed.

Abbreviations. HIV, Human Immunodeficiency virus. KO, knock out. R5, CCR5. X4, CXCR4. CAR, Chimeric Antigen Receptor. bPD1-CAR, blocking anti-PD-1 CAR. nbPD1-CAR, non-blocking anti-PD-1 CAR. VL, viral load. ACT, adoptive cell transfer. ART, antiretroviral therapy. BM, bone marrow.

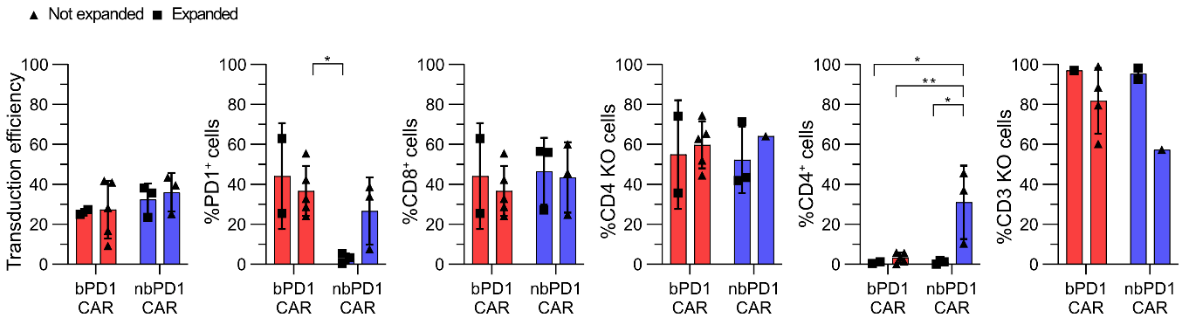

**Fig. S8. CAR-T cells phenotype before *in vivo* infusion**

Phenotype of CAR-T cells before injection in HIV infected hu-mice comparing the ones that expanded *in vivo* versus the ones that did not. Mean ± SD of 3 to 5 donors is shown. Two-way ANOVA, Tukey's multiple comparisons test.

<u>Abbreviations</u>. KO, knock out. CAR, Chimeric Antigen Receptor. bPD1-CAR, blocking anti-PD-1 CAR. nbPD1-CAR, non-blocking anti-PD-1 CAR.

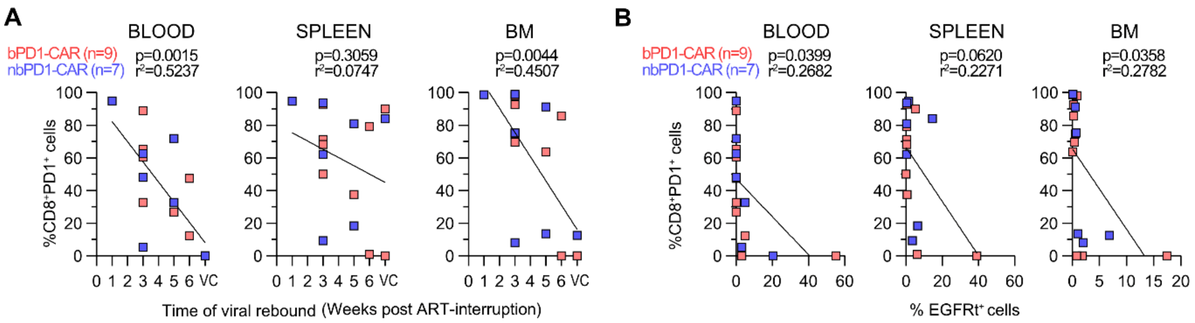

**Fig. S9. Correlations between the number of CD8$^+$PD-1$^+$ cells, viral rebound and CAR-T detection**

(A) Correlation between CD8$^+$PD-1$^+$ cells (gated in huCD45$^+$EGFRt$^-$ cells) and the time of viral rebound in the ACT-on-ART group (bPD1-CAR, n=9, nbPD1-CAR, n=7). Simple linear regression was used.

(B) Correlation between CD8$^+$PD-1$^+$ cells (gated in huCD45$^+$EGFRt$^-$ cells) and CAR-T cells detection in the ACT-on-ART group (bPD1-CAR, n=9, nbPD1-CAR, n=7). Simple linear regression was used.

Abbreviations. CAR, Chimeric Antigen Receptor. bPD1-CAR, blocking anti-PD-1 CAR. nbPD1-CAR, non-blocking anti-PD-1 CAR. VC, viral control. EGFRt, Truncated epidermal growth factor receptor. ART, antiretroviral therapy. BM, bone marrow.

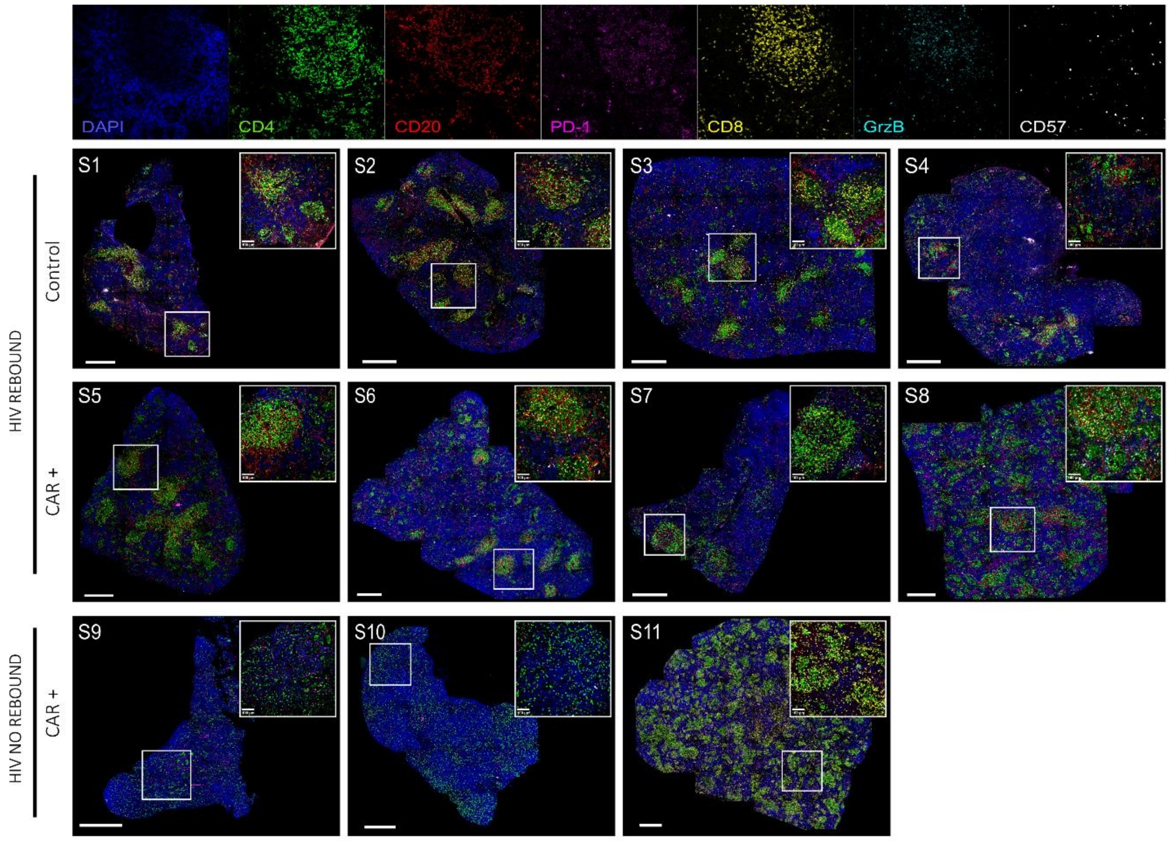

**Fig. S10. Confocal images of full tissue sections of spleens from humanized mice**

Overview images of the full spleen sections acquired at the confocal microscope (20x magnification). Areas selected for higher resolution images of figure 6 are shown. Immunofluorescence multicolour panel showing DAPI (blue), CD4 (green), CD20 (red), PD-1 (magenta), CD8 (yellow), GrzB (cyan) and CD57 (gray). Scale bar, 500 µm.
Abbreviations. HIV, Human Immunodeficiency virus. CAR, Chimeric Antigen Receptor.

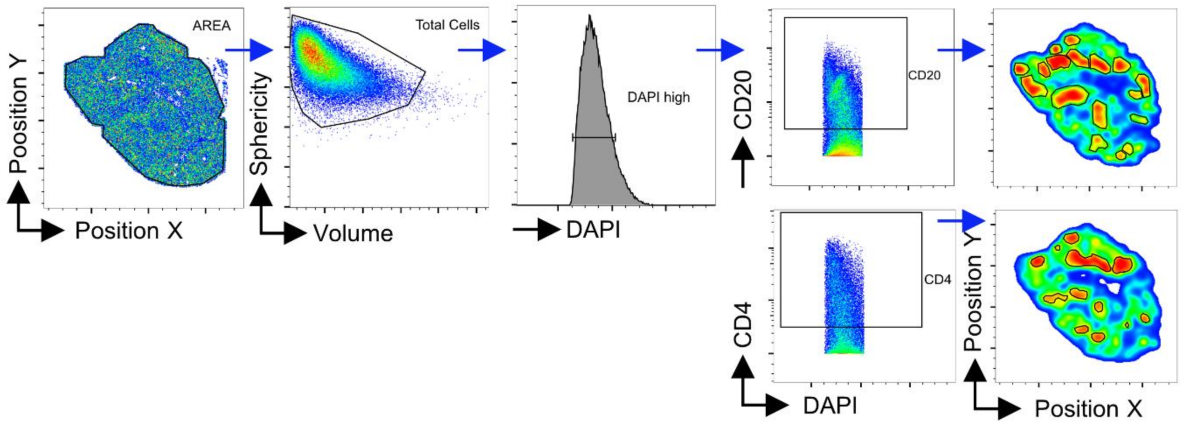

**Fig. S11. Gating strategy for histo-cytometry analysis**

Gating strategy for histo-cytometry analysis of CD20 or CD4-enriched zones, based on cell density.

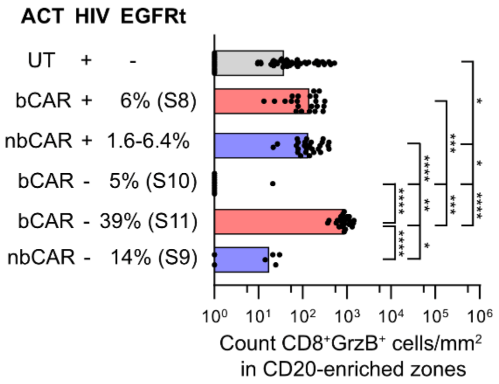

**Fig. S12. Histo-cytometry quantification of $CD8^{+}GrzB^{+}$ cells in CD20-enriched zones**
Histo-cytometry analysis showing $CD8^{+}$ $GrzB^{+}$ cell counts/$mm^2$ in CD20-enriched zones. Each dot represents a CD20-enriched zones (n=155). All 13 mice shown in Fig.6B are included. Median is shown. Kruskal-Wallis test and Dunn's test.

Abbreviations. bCAR, blocking anti-PD-1 CAR. nbCAR, non-blocking anti-PD-1 CAR. GrzB, Granzyme B. EGFRt, Truncated epidermal growth factor receptor. UT, untransduced T cells.

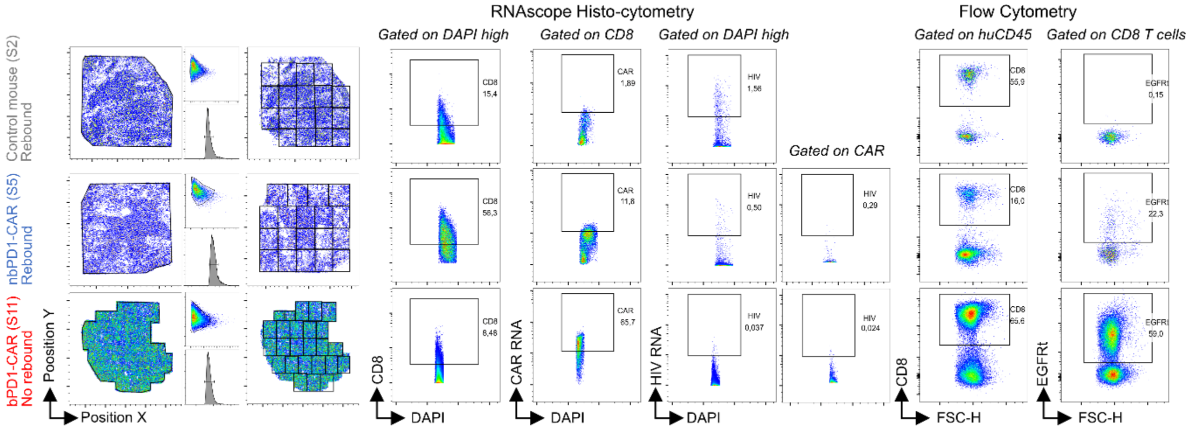

**Fig. S13. RNAscope histo-cytometry gating strategy**

Gating strategy for histo-cytometry analysis of RNAscope data (left). RNAscope histo-cytometry and flow cytometry comparison of mice S2, S5, S11 (right). HIV RNA$^{+}$ cells gated on DAPI, CAR$^{+}$ total T cells and CAR$^{+}$ non-CD8 T cells.

Abbreviations. bCAR, blocking anti-PD-1 CAR. nbCAR, non-blocking anti-PD-1 CAR. GrzB, Granzyme B. EGFRt, Truncated epidermal growth factor receptor.

### Supplementary tables

**Table S1.** Viremia levels pre- and post- CAR-T cells treatment, mice death rates and huCD45 and EGFRt flow cytometry levels in organs.

| Viremia before ART | MOUSE ID | blood (1 week post ART interruption - 3 | blood (3 weeks post ART interruption - 5 | blood (5 weeks post ART interruption - 7 | blood (6/8 weeks post ART interruption - 8/10 weeks post ACT) | EGFR detection in organs (% of huCD45) | | | % of huCD45+ cells in organs | | |
|---|---|---|---|---|---|---|---|---|---|---|---|
| UT T cells | | | | | | SPLEEN | BM | BLOOD | SPLEEN | BM | BLOOD |
| 1.22E+05 | 5629 | NEG | 5.03E+05 | *missing* | 1.02E+04 | | | | 20.4 | 0.66 | 2.04 |
| 1.46E+04 | 5630 | NEG | 2.76E+04 | *dead* | *dead* | | | | | | |
| 7.83E+03 | 5660 | NEG | 7.87E+03 | *missing* | 5.42E+04 | | | | 59 | 18 | 25.5 |
| 1.50E+05 | 5674 | NEG | 1.43E+03 | *missing* | 4.58E+02 | | | | 11.8 | 0.1 | 0.1 |
| 3.87E+03 | 5659 | NEG | 2.69E+04 | *missing* | 2.55E+05 | | | | 26.4 | 47.3 | 59.5 |
| 7.12E+04 | 5693 | NEG | 1.21E+04 | 7.11E+04 | *dead* | | | | | | |
| 1.77E+04 | 5720 | NEG | 7.16E+04 | 2.02E+05 | *dead* | | | | | | |
| 2.21E+04 | 5727 | NEG | 5.23E+04 | 2.96E+04 | 2.45E+04 | | | | 24 | 3.99 | 7.98 |
| 2.29E+03 | 5730 | NEG | 1.97E+04 | 8.78E+04 | 1.34E+05 | | | | 85.2 | 71.7 | 2.99 |
| bPD1-CAR | | | | | | | | | | | |
| 8.97E+03 | 5672 | 1.47E+03 | *dead* | *dead* | *dead* | | | | | | |
| 3.87E+03 | 5641 | NEG | 1.27E+04 | *missing* | 4.26E+04 | 0.19 | 0.21 | 0.23 | 40.6 | 4.11 | 9.48 |
| 7.22E+04 | 5657 | NEG | 1.51E+04 | *missing* | 3.08E+05 | 0.19 | 0.87 | 0.00 | 25.3 | 1.91 | 4.23 |
| 5.30E+04 | 5658 | NEG | 1.57E+03 | *missing* | 1.88E+05 | 0.24 | 0.41 | 0.04 | 74.9 | 35.6 | 30.1 |
| 5.85E+04 | 5722 | NEG | 4.45E+03 | *dead* | *dead* | | | | | | |
| 1.08E+04 | 5723 | NEG | NEG | NEG | *dead* | | | | | | |
| 1.07E+05 | 5750 | NEG | NEG | NEG | *dead* | | | | | | |
| 7.09E+03 | 5754 | NEG | NEG | 1.04E+04 | 9.72E+03 | 0.59 | 0.087 | 0.17 | 22.2 | 74 | 30.9 |
| 8.91E+03 | 5675 | NEG | *missing* | *missing* | NEG | 5.19 | 17.41 | 3.12 | 13.6 | 0.71 | 1.06 |
| 1.27E+04 | 5733 | NEG | NEG | NEG | NEG | 39.2 | 1.92 | 55.2 | 39.8 | 2.09 | 79.8 |
| 2.81E+03 | 5734 | NEG | NEG | NEG | 1.49E+03 | 6.24 | 0.81 | 5 | 64.3 | 52.4 | 30.6 |
| 3.14E+03 | 5882 | NEG | 6.69E+05 | *missing* | 4.77E+06 | 0.178 | 0.59 | 0.027 | 52.3 | 48 | 1.94 |
| 3.24E+03 | 5886 | NEG | NEG | *missing* | 1.15E+05 | 0.218 | 0.3 | 0.012 | 49.9 | 49 | 25.4 |
| nbPD1-CAR | | | | | | | | | | | |
| 1.62E+05 | 5669 | NEG | 1.40E+03 | *dead* | *dead* | | | | | | |
| 7.18E+02 | 5644 | NEG | NEG | 7.30E+03 | 3.34E+04 | 0.19 | 0.56 | 0.06 | 64.2 | 18.9 | 22.7 |
| 1.24E+04 | 5645 | NEG | 6.20E+02 | *dead* | *dead* | | | | | | |
| 1.73E+03 | 5681 | NEG | 7.96E+02 | 2.57E+04 | 3.15E+05 | 0.54 | 0.18 | 0.00 | 35.1 | 17 | 5.43 |
| 1.10E+03 | 5724 | NEG | NEG | NEG | *dead* | | | | | | |
| 1.12E+04 | 5736 | NEG | 4.73E+04 | 1.01E+05 | 6.02E+04 | 3.51 | 2.05 | 3.1 | 39.1 | 26.9 | 29 |
| 9.95E+02 | 5737 | NEG | NEG | 2.41E+04 | 3.54E+04 | 6.4 | 1.06 | 5.01 | 35.6 | 41.5 | 26.7 |
| 1.02E+04 | 5679 | 2.83E+03 | *missing* | *missing* | 3.03E+05 | 1.58 | 0.14 | 0.00 | 70.7 | 10.4 | 4.87 |
| 1.94E+03 | 5676 | NEG | *missing* | *missing* | NEG | 14.57 | 6.90 | 20.50 | 22.4 | 61.3 | 51.2 |
| 1.05E+03 | 5888 | NEG | 2.01E+06 | *missing* | 1.90E+05 | 0.189 | 0.669 | 0.026 | 54.2 | 47.4 | 15.3 |
| 1.80E+03 | 5885 | NEG | NEG | *missing* | *dead* | | | | | | |

**Table S2.** Subgrouping of the mice in CAR detectable and non-detectable groups based on 1% threshold of $huCD45^{+}EGFRt^{+}$ cells.

EGFRt Flow Cytometry frequencies

| | CAR detectable | | | | CAR non detectable | | |
|---|---|---|---|---|---|---|---|
| bPD1-CAR | | | | | | | |
| MOUSE | **SPLEEN** | **BM** | **BLOOD** | MOUSE | **SPLEEN** | **BM** | **BLOOD** |
| 5675 | 5.19 | 17.41 | 3.12 | 5754 | 0.59 | 0.087 | 0.17 |
| 5733 | 39.2 | 1.92 | 55.2 | 5641 | 0.19 | 0.21 | 0.23 |
| 5734 | 6.24 | 0.81 | 5 | 5657 | 0.19 | 0.87 | 0.00 |
| | | | | 5658 | 0.24 | 0.41 | 0.04 |
| | | | | 5882 | 0.178 | 0.59 | 0.027 |
| | | | | 5886 | 0.218 | 0.3 | 0.012 |
| nbPD1-CAR | | | | | | | |
| 5676 | 14.57 | 6.90 | 20.50 | 5644 | 0.19 | 0.56 | 0.06 |
| 5679 | 1.58 | 0.14 | 0.00 | 5681 | 0.54 | 0.18 | 0.00 |
| 5736 | 3.51 | 2.05 | 3.1 | 5888 | 0.189 | 0.669 | 0.026 |
| 5737 | 6.4 | 1.06 | 5.01 | | | | |

**Table S3.** Tables show all different factors involved in CAR-T cells generation and ACT, comparing the mice groups based on CAR detection or viral rebound.

| | no HIV rebound N =3[1] | HIV rebound N = 13[1] | p-value[2] |
|---|---|---|---|
| **Mice with detectable CAR** | 3 / 3 (100%) | 4 / 13 (31%) | 0.063 |
| **Mouse sex** | | | **0.036** |
| F | 3 / 3 (100%) | 3 / 13 (23%) | |
| M | 0 / 3 (0%) | 10 / 13 (77%) | |
| **Mice cohort** | | | >0.9 |
| 1 | 2 / 3 (67%) | 6 / 13 (46%) | |
| 2 | **1 / 3 (33%)** | 4 / 13 (31%) | |
| 3 | 0 / 3 (0%) | 3 / 13 (23%) | |
| **Mouse donor** | | | 0.5 |
| T874 | 0 / 3 (0%) | 1 / 13 (7.7%) | |
| T875 | 0 / 3 (0%) | 1 / 13 (7.7%) | |
| HFL CD34+ Dec 2016 | 0 / 3 (0%) | 1 / 13 (7.7%) | |
| HFL CD34+ Oct2020 | 0 / 3 (0%) | 1 / 13 (7.7%) | |
| T787 | 0 / 3 (0%) | 1 / 13 (7.7%) | |
| T794 | 2 / 3 (67%) | 0 / 13 (0%) | |
| T814 | 0 / 3 (0%) | 2 / 13 (15%) | |
| T849 | 1 / 3 (33%) | 3 / 13 (23%) | |
| T874 | 0 / 3 (0%) | 1 / 13 (7.7%) | |
| U320 | 0 / 3 (0%) | 2 / 13 (15%) | |
| **Cells origin** | | | >0.9 |
| CD34- | 2 / 3 (67%) | 7 / 13 (54%) | |
| spleen | 1 / 3 (33%) | 6 / 13 (46%) | |
| **Type of CAR** | | | >0.9 |
| bPD1 | 2 / 3 (67%) | 7 / 13 (54%) | |
| nbPD1 | 1 / 3 (33%) | 6 / 13 (46%) | |
| **Number of total injected cells (MIO)** | 3.20 [1.96, 8.00] | 4.00 [3.00, 8.00] | 0.5 |
| **Number of injected CAR T cells** | 1.20 [0.55, 2.00] | 1.00 [0.80, 1.50] | >0.9 |
| **CRISPR editing** | | | 0.8 |
| CD3/CD4/PD1 KO | 0 / 3 (0%) | 4 / 13 (31%) | |
| CD4/CD3/PD1 | 2 / 3 (67%) | 3 / 13 (23%) | |
| CD4/PD1 | 1 / 3 (33%) | 4 / 13 (31%) | |
| Unedited | 0 / 3 (0%) | 2 / 13 (15%) | |
| **% CD4 T cells** | 1 [0, 1] | 3 [1, 9] | 0.092 |
| **% CD4 KO T cells** | 72 [36, 74] | 45 [42, 68] | 0.4 |
| **% CD8 T cells** | 27 [26, 63] | 46 [24, 56] | >0.9 |
| **Ratio CD8/CD4** | 0.37 [0.34, 1.70] | 1.00 [0.32, 1.27] | >0.9 |
| **State of cells at ACT** | | | 0.2 |
| fresh | 1 / 3 (33%) | 10 / 13 (77%) | |
| frozen | 2 / 3 (67%) | 3 / 13 (23%) | |
| **Duration of expansion (days)** | 12.00 [11.00, 12.00] | 11.00 [11.00, 11.00] | 0.2 |
| **Engraftment before ACT (%huCD45)** | 23 [23, 64] | 29 [20, 52] | 0.8 |
| **Plasma VL before ACT** | 8,910 [1,940, 12,700] | 3,240 [1,730, 10,200] | 0.6 |
| **Plasma VL at sacrifice** | 0 [0, 0] | [illegible]5,000 [35,400, 303,0[illegible] | **0.010** |
| **Engraftment at sacrifice (%huCD45 blood)** | 51 [1, 80] | 23 [5, 29] | 0.4 |

[1] n / N (%); Median [IQR]
[2] Fisher's exact test; Wilcoxon rank sum test; Wilcoxon rank sum exact test

| | no detectable CAR N = 9[1] | detectable CAR N = 7[1] | p-value[2] |
|---|---|---|---|
| **HIV rebound** | 9 / 9 (100%) | 4 / 7 (57%) | 0.063 |
| **Mouse sex** | | | 0.3 |
| F | 2 / 9 (22%) | 4 / 7 (57%) | |
| M | 7 / 9 (78%) | 3 / 7 (43%) | |
| **Mice cohort** | | | 0.14 |
| 1 | 5 / 9 (56%) | 3 / 7 (43%) | |
| 2 | 1 / 9 (11%) | 4 / 7 (57%) | |
| 3 | 3 / 9 (33%) | 0 / 7 (0%) | |
| **Mouse donor** | | | **0.044** |
| T874 | 1 / 9 (11%) | 0 / 7 (0%) | |
| T875 | 1 / 9 (11%) | 0 / 7 (0%) | |
| HFL CD34+ Dec 2016 | 1 / 9 (11%) | 0 / 7 (0%) | |
| HFL CD34+ Oct2020 | 1 / 9 (11%) | 0 / 7 (0%) | |
| T787 | 1 / 9 (11%) | 0 / 7 (0%) | |
| T794 | 0 / 9 (0%) | 2 / 7 (29%) | |
| T814 | 1 / 9 (11%) | 1 / 7 (14%) | |
| T849 | 0 / 9 (0%) | 4 / 7 (57%) | |
| T874 | 1 / 9 (11%) | 0 / 7 (0%) | |
| U320 | 2 / 9 (22%) | 0 / 7 (0%) | |
| **Cells origin** | | | 0.6 |
| CD34- | 6 / 9 (67%) | 3 / 7 (43%) | |
| spleen | 3 / 9 (33%) | 4 / 7 (57%) | |
| **Type of CAR** | | | 0.6 |
| bPD1 | 6 / 9 (67%) | 3 / 7 (43%) | |
| nbPD1 | 3 / 9 (33%) | 4 / 7 (57%) | |
| **Number of total injected cells (MIO)** | 3.90 [3.00, 4.50] | 8.00 [3.20, 8.00] | 0.4 |
| **Number of injected CAR T cells** | 0.90 [0.80, 1.00] | 2.00 [1.20, 2.00] | **0.040** |
| **CRISPR editing** | | | 0.056 |
| CD3/CD4/PD1 KO | 4 / 9 (44%) | 0 / 7 (0%) | |
| CD4/CD3/PD1 | 2 / 9 (22%) | 3 / 7 (43%) | |
| CD4/PD1 | 1 / 9 (11%) | 4 / 7 (57%) | |
| Unedited | 2 / 9 (22%) | 0 / 7 (0%) | |
| **% CD4 T cells** | 9 [3, 12] | 1 [0, 2] | **0.020** |
| **% CD4 KO T cells** | 63 [45, 70] | 42 [36, 72] | 0.6 |
| **% CD8 T cells** | 33 [22, 46] | 56 [27, 63] | 0.056 |
| **Ratio CD8/CD4** | 0.50 [0.28, 1.00] | 1.27 [0.37, 1.70] | 0.056 |
| **State of cells at ACT** | | | 0.6 |
| fresh | 7 / 9 (78%) | 4 / 7 (57%) | |
| frozen | 2 / 9 (22%) | 3 / 7 (43%) | |
| **Duration of expansion (days)** | 11.00 [11.00, 11.00] | 11.00 [11.00, 12.00] | 0.4 |
| **Engraftment before ACT (%huCD45)** | 36 [16, 60] | 26 [23, 39] | >0.9 |
| **Plasma VL before ACT** | 3,240 [1,730, 7,090] | 8,910 [1,940, 11,200] | 0.8 |
| **Plasma VL at sacrifice** | [illegible]8,000 [42,600, 308,00[illegible] | 1,490 [0, 60,200] | **0.034** |
| **Engraftment at sacrifice (%huCD45 blood)** | 15 [5, 25] | 29 [5, 51] | 0.3 |

[1] n / N (%); Median [IQR]
[2] Fisher's exact test; Wilcoxon rank sum test; Wilcoxon rank sum exact test

**Table S4.** Antibodies used for flow cytometry.

| Target | Clone | Fluorophore | Vendor | Catalog number | Country |
|---|---|---|---|---|---|
| CD3 | UCHT-1 | PE-Cy7 | BD Biosciences | 563423 | Franklin Lakes, NJ, USA |
| CD3 | OKT3 | BV785 | BioLegend | 317330 | San Diego, CA, USA |
| CD4 | RPA-T4 | AF700 | BD Biosciences | 557922 | Franklin Lakes, NJ, USA |
| CD4 | RPA-T4 | FITC | BD Biosciences | 555346 | Franklin Lakes, NJ, USA |
| CD4 | SK3 | BUV395 | BD Biosciences | 563550 | Franklin Lakes, NJ, USA |
| CD4 | RPA-T4 | PE-Cy7 | BioLegend | 300512 | San Diego, CA, USA |
| CD8 | SK1 | BV421 | BioLegend | 344748 | San Diego, CA, USA |
| CD8 | RPA-T8 | APC-Cy7 | BD Biosciences | 557760 | Franklin Lakes, NJ, USA |
| CD8 | SK1 | PE | BD Biosciences | 345773 | Franklin Lakes, NJ, USA |
| CD25 | CD25-4E3 | APC | Thermo Fisher Scientific | 17-0257-42 | Waltham, MA, USA |
| CD71 | CY1G4 | APC-Cy7 | BioLegend | 334110 | San Diego, CA, USA |
| CD71 | M-A712 | FITC | BD Biosciences | 555536 | San Diego, CA, USA |
| CTV | CellTrace Violet | Pacific Blue | Invitrogen | C34557 | Waltham, MA, USA |
| CD45 | H130 | FITC | BioLegend | 304006 | San Diego, CA, USA |
| CD45RA | HI100 | BV650 | BD Biosciences | 563963 | Franklin Lakes, NJ, USA |
| CD62L | DREG-56 | BV-421 | BD Biosciences | 563862 | Franklin Lakes, NJ, USA |
| LAG3 | 11C3C65 | PerCP/Cyanine5.5 | BioLegend | 369312 | San Diego, CA, USA |
| TIM3 | CF38-2E2 | BV 785 | BioLegend | 345032 | San Diego, CA, USA |
| PD1 | EH12.2H7 | APC | BioLegend | 329908 | San Diego, CA, USA |
| PD-1 | EH12.1 | PE-Cy7 | BD Biosciences | 561272 | Franklin Lakes, NJ, USA |
| CCR5 | REA245 | APC | Miltenyi Biotec GmbH | 130-120-057 | Miltenyi Biotec, Bergisch-Gladbach, Germany |
| CXCR4 | 12G5 | PE-Cy7 | BioLegend | 306514 | San Diego, CA, USA |
| EGFR | AY13 | PE | BioLegend | 352904 | San Diego, CA, USA |
| EGFR | AY13 | FITC | BioLegend | 352908 | San Diego, CA, USA |
| DAPI | Live/Dead | NA | Invitrogen | D1306 | Waltham, MA, USA |
| Anti-his tag | J095G46 | PE | BioLegend | 362603 | San Diego, CA, USA |
| Streptavidin | | PE | BD Biosciences | 554061 | Franklin Lakes, NJ, USA |
| Goat anti-Human IgG (H+L) Cross-Adsorbed Secondar | | Alexa Fluor 568 | Invitrogen | A-21090 | Waltham, MA, USA |
| LIVE/DEAD® Fixable Aqua Dead Cell Stain Kit | | NA | Thermo Fisher Scientific | L34957 | Waltham, MA, USA |
| Zombie NIR Fixable Viability Kit | | NA | BioLegend | 423105/423106 | San Diego, CA, USA |

**Table S5.** crRNA sequences.

| Name | Sequence |
|---|---|
| TRBC | CCCACCAGCTCAGCTCCACG |
| CD4 | GGCAAGGCCACAATGAACCG |
| PD-1 | CTGCAGCTTCTCCAACACAT |
| HLA-A2 | CCTCGTCCTGCTACTCTCGG |
| CCR5 | CAATGTGTCAACTCTTGACA |
| CXCR4 | CACTTCAGATAACTACACCG |

**Table S6.** Antibodies used for immunofluorescence staining of spleen tissues.

| Target | Clone | Dilution | Catalogue Number | Fluorophore | Vendor |
|---|---|---|---|---|---|
| CD4 | EPR6855 | 1/300 | ab133616 | | Abcam |
| PD1 | NAT105 | 1/100 | 3137 | | Bio Optica |
| CD57 | NK-1 | 1/200 | Mob163 | | CliniSciences |
| CD20 | L26 | 1/400 | NCL-L-CD20-L26 | | Leica system |
| CD8 | C8/144b | 1/50 | M7103 | | Agilent |
| GrzB | GrB-7 | 1/40 | MON7029C | | Monosan |
| CD4 | Polyclonal | 1/35 | FAB8165N | AF 700 | Bio-techne |
| CD8 | C8/144B | 1/25 | 372906 | AF 647 | Biolegend |
